# Pioneer-factor activity requires stable chromatin occupancy mediated by both sequence-specific binding and disordered protein domains

**DOI:** 10.64898/2026.02.06.703578

**Authors:** Meghan M. Freund, F. Javier deHaro-Arbona, Sarah Baloul, Ali Torhorst, Charalambos Roussos, Abby J. Ruffridge, Andrew Q. Rashoff, Ryen Hazzard, Peter W. Lewis, Sarah J. Bray, Melissa M. Harrison

## Abstract

Pioneer transcription factors overcome the restrictive barrier imposed by chromatin to drive cell-fate specification, yet how their domains collectively support this activity remains unclear. Here, we use the deeply conserved pioneer factor Grainy head to define the protein-intrinsic features that govern pioneering activity. By integrating biochemistry, genomics and quantitative live-cell imaging, we determined that both the conserved DNA-binding domain and the extended, intrinsically disordered N-terminus are required for the stable chromatin occupancy that supports access to closed chromatin and the induction of chromatin accessibility. The disordered N-terminus supports pioneer activity through interactions that do not rely on strict amino acid sequence but instead overall composition. While our results show that pioneering activity depends on the combinatorial contributions of structured and disordered domains, mitotic retention depends solely on sequence-specific DNA binding. These results support stable chromatin occupancy mediated by multiple protein domains as necessary for pioneering function and that this is separable from the mechanisms required for mitotic retention.

## INTRODUCTION

The gene-expression programs that control cellular identity are driven by transcription factors that recognize specific DNA sequence motifs. However, transcription-factor binding is constrained by the packaging of DNA into chromatin, which restricts access to much of the genome (*1*). A subset of transcription factors, termed pioneer factors, overcome this barrier by binding to nucleosomal DNA, promoting local chromatin accessibility, and enabling broad transcriptional reprogramming events (*2–5*). These features allow pioneer factors to function as drivers of cell-fate specification. A set of characteristics has been used to define pioneer factors, including binding nucleosomes *in vitro,* binding closed, nucleosome-occupied chromatin in cells, promoting chromatin accessibility, and being retained on mitotic chromosomes (*6*). Nonetheless, these properties are not shared by all pioneer factors, and the mechanistic details by which individual factors engage and open chromatin differ (*6*). To address how pioneer factors uniquely bind the genome, it is necessary to systematically test the relationship between these defining properties.

Structured DNA-binding domains (DBDs) provide sequence specificity for transcription factors and, in many cases, are sufficient for nucleosome binding by pioneer factors *in vitro* (*7–11*). Yet, recent *in vivo* assays demonstrate that DBDs alone cannot explain the full spectrum of pioneering activity (*12*, *13*). Indeed, non-DBDs contribute to chromatin engagement (*14–19*) and in some cases can direct binding on their own (*15*). These data demonstrate that eukaryotic transcription factors are not as modular as their prokaryotic counterparts. Pioneer factors contain diverse non-DBDs that differ widely in composition, yet frequently include intrinsically disordered regions (IDRs). These IDRs have been proposed to facilitate target search, stabilize chromatin interactions, recruit cofactors, and drive condensate formation (*19–27*). Despite the emerging roles of IDRs, they lack conserved sequence motifs, making the mechanism by which they support pioneering unclear.

Grainy head (Grh) is an essential and deeply conserved transcription factor that drives expression of genes that promote epithelial cell fate in organisms ranging from worms to humans (*28–37*). Grh was initially identified in *Drosophila melanogaster*, where it is encoded by a single gene (*28*, *38–40*). By contrast, mammals have three Grainyhead-like proteins (GRHL1-3) whose dysregulation is associated with numerous epithelial cancers (*32*, *34*, *41–43*). Grh proteins from both flies and mice exhibit key features of pioneer factors: loss of Grh reduces chromatin accessibility, whereas ectopic expression promotes accessibility at previously closed sites (*12*, *44–46*). The structure of Grh is shared across species and contains a highly conserved DBD and dimerization domain (*30*, *32*, *47*). In all organisms studied, Grh binds the same DNA motif, reflecting the strong conservation of the Grh DBD (*31*, *34*, *35*). By contrast, the N-terminus is poorly conserved and shows striking divergence in length, sequence and predicted structure (*48*). Because of the shared roles of GRHL proteins in pioneering and their impact on human health, we use the simplified *Drosophila* system to determine how both ordered and disordered protein domains individually contribute to multiple, defining features of pioneer factors. By integrating biochemical, genomic and imaging assays, we reveal that the stable chromatin occupancy required for pioneering activity depends on sequence-specific DNA binding and additional protein domains, which can vary in sequence and structure. By contrast, mitotic retention relies solely on sequence-specific DNA binding.

## RESULTS

### Grh has features of a pioneer transcription factor

To discover how distinct features of pioneer factors relate to each other, we first determined the essential pioneering features of Grh. Despite being identified as a pioneer factor, whether Grh can directly engage nucleosomal DNA has not been tested directly (*12*, *44–46*). Therefore, we reconstituted nucleosomes containing an endogenous target sequence and tested binding by full-length recombinant Grh *in vitro*. We identified a genomic locus in closed chromatin that contains a canonical Grh-binding motif and becomes accessible upon Grh expression and binding in *Drosophila* Schneider 2 (S2) cells (Figure S1A) (*12*). We generated nucleosomes using a 159 bp fragment surrounding the Grh motif. In electrophoretic mobility shift assays (EMSAs), full-length recombinant Grh bound to both free and nucleosomal DNA (Figure 1A). Together, these data establish that Grh directly interacts with nucleosomal DNA, a defining feature of pioneer factors.

**Figure 1.**
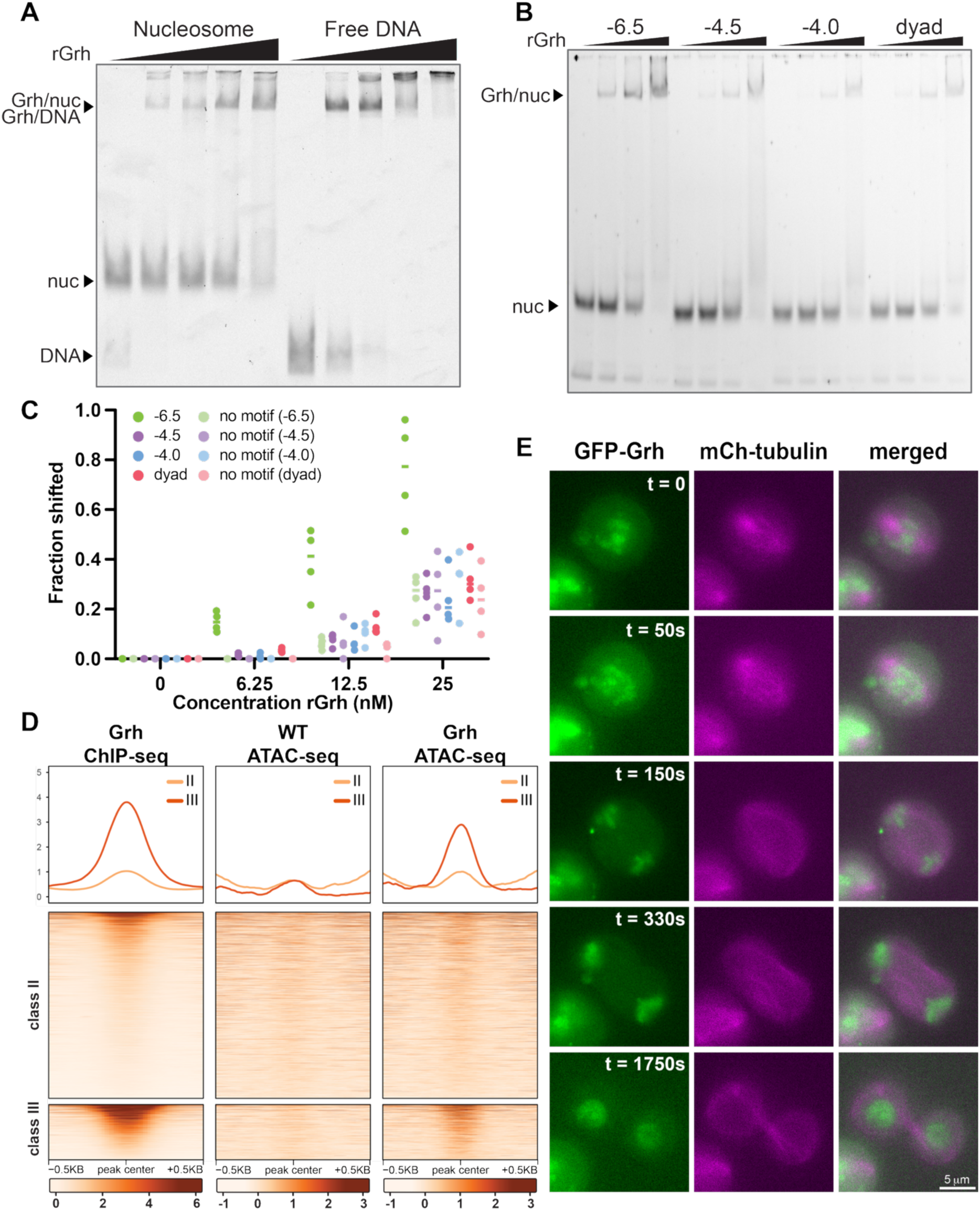
Grainy head has features of a pioneer transcription factor. (**A**) EMSA with increasing concentrations of recombinant Grh (rGrh) (0 – 50 nM) incubated with Cy5-labeled free or nucleosomal DNA. (**B**) EMSA with increasing concentrations of rGrh (0 – 25nM) incubated with Cy5-labeled Widom 601 nucleosomes with the canonical Grh motif positioned at SHLs -6.5, -4.5, -4.0 or 0 (dyad). (**C**) Quantification of the fraction of Grh-bound nucleosomes, indicated by the proportion shifted in the EMSA shown in (B) and three additional experiments. (**D**) Heatmaps and metaplots of binding by Grh (ChIP-seq) and chromatin accessibility (ATAC-seq) in wild-type (WT) S2 cells or those in which Grh is induced. Classes from Gibson et al. 2024 (*12*). (**E**) Images of S2 cells expressing GFP-Grh (green) and mCherry-tubulin (magenta) over mitosis. t = time in seconds relative to the initial image (metaphase).

Pioneer factors vary in their recognition of nucleosomal motifs, with some binding at the dyad, some at the edge, and others at sites in between (*10*, *49*, *50*). To test the motif position preference for Grh, we took advantage of the well-positioned Widom 601 nucleosome, allowing us to insert the Grh motif at defined positions around the nucleosome axis (*51*). We placed the motif at the edge (super helical location (SHL) -6.5), internal positions (SHL -4.5 and -4.0 solvent exposed and buried, respectively), and the dyad (SHL 0) and assayed binding by recombinant Grh (Fig. S1B). Motif-specific binding was observed when the Grh motif was positioned at SHL - 6.5, whereas binding at internal and central positions was not enriched above background (Fig. 1B, C, Fig. S1C). This preference resembles that reported for other pioneer factors and likely reflects the dynamic nature of DNA wrapping at the nucleosome entry/exit site (*49*, *52*, *53*).

To examine pioneering activity in a cellular context, we ectopically expressed Grh at physiological levels in S2 cells, which do not endogenously express Grh and assayed genome-wide binding using chromatin immunoprecipitation coupled with sequencing (ChIP-seq) and chromatin accessibility using assay for transposase-accessible chromatin coupled with sequencing (ATAC-seq) (*12*). As reported previously, Grh-bound sites fall into three categories distinguished by their accessibility before and after Grh binding (*12*). Class I sites are accessible regions that Grh occupies without altering accessibility and are sites of promiscuous, non-specific binding (Fig. S2) (*12*). Class II and class III sites are inaccessible prior to binding and reflect the ability of Grh to function as a pioneer factor. These classes differ in their response to Grh binding with class II sites remaining inaccessible and class III sites becoming accessible (Fig. 1D, Fig. S2). These data demonstrate that at a subset of loci Grh functions as a pioneer factor by binding to closed chromatin and promoting accessibility.

Another feature commonly associated with pioneer factors is mitotic retention. During mitosis, most transcription factors are not retained on chromatin with only a subset remaining bound (mitotically retained) (*54–56*). These mitotically retained factors are thought to facilitate rapid reactivation of gene expression after mitosis (*57–61*).

Although the mechanisms of mitotic retention are not well understood, features that promote pioneer activity are thought to contribute to retention, as many pioneer factors are mitotically retained (*45*, *57–59*, *61–66*). Indeed, endogenous Grh is mitotically retained at euchromatin in the gastrulating embryo (*45*). We examined whether this feature is shared in S2 cells, a tractable cellular system. Using mCherry-tubulin to mark mitotic cells, we identified a clear association of transiently expressed GFP-Grh with mitotic chromosomes, recapitulating prior *in vivo* observations (Fig. 1E) (*45*). These results, along with previous data, demonstrate that Grh possesses multiple, defining features of pioneer factors and establish assays to test the contribution of various protein domains to these pioneering activities (*12*, *44–46*).

### The DNA-binding domain of Grh and regions outside are required for nucleosome binding and opening chromatin

Having established the core pioneering features of Grh, we next determined how the DNA-binding domain and the disordered N-terminus contributed to *in vitro* nucleosome binding and how this relates to the ability of Grh to promote chromatin accessibility in culture. To test the role of DNA binding, we leveraged the high degree of conservation in the DNA-binding domain (DBD) and the published structure of the mammalian Grainy head like protein GRHL1 to identify residues that directly contact DNA (*67*). We generated purified full-length protein with mutations that are predicted to abrogate binding to either the phosphate backbone (Grh^C800A,K807A^) or the conserved guanine in the Grh motif (Grh^R806A^). Consistent with the role of these residues in promoting DNA binding, both DBD mutations reduced binding to free DNA, with Grh^R806A^ showing a stronger decrease than Grh^C800A,K807A^ (Fig. 2A, B, Fig. S3A). To assess the influence of the large, disordered N-terminus, we purified protein that corresponded to only the C-terminal DBD and dimerization domains (Grh^ΔN^). Grh^ΔN^ had decreased binding to free DNA relative to wild-type protein but retained some DNA-binding activity (Fig. 2A, B, Fig. S3A). Because pioneering involves interactions with DNA that is wrapped around histones, we tested the capacity of each mutant protein to bind nucleosomes. Similar to binding to free DNA, both DBD mutants reduced nucleosome binding, with Grh^R806A^ showing the strongest decrease, and Grh^ΔN^ exhibiting a modest but statistically significant reduction relative to wild-type Grh (Fig. 2C, D). These results indicate that *in vitro* nucleosome binding reflects the ability to bind free DNA and requires both the DBD and N-terminal domain, with a particular dependence on R806, which mediates sequence-specific DNA contacts.

**Figure 2.**
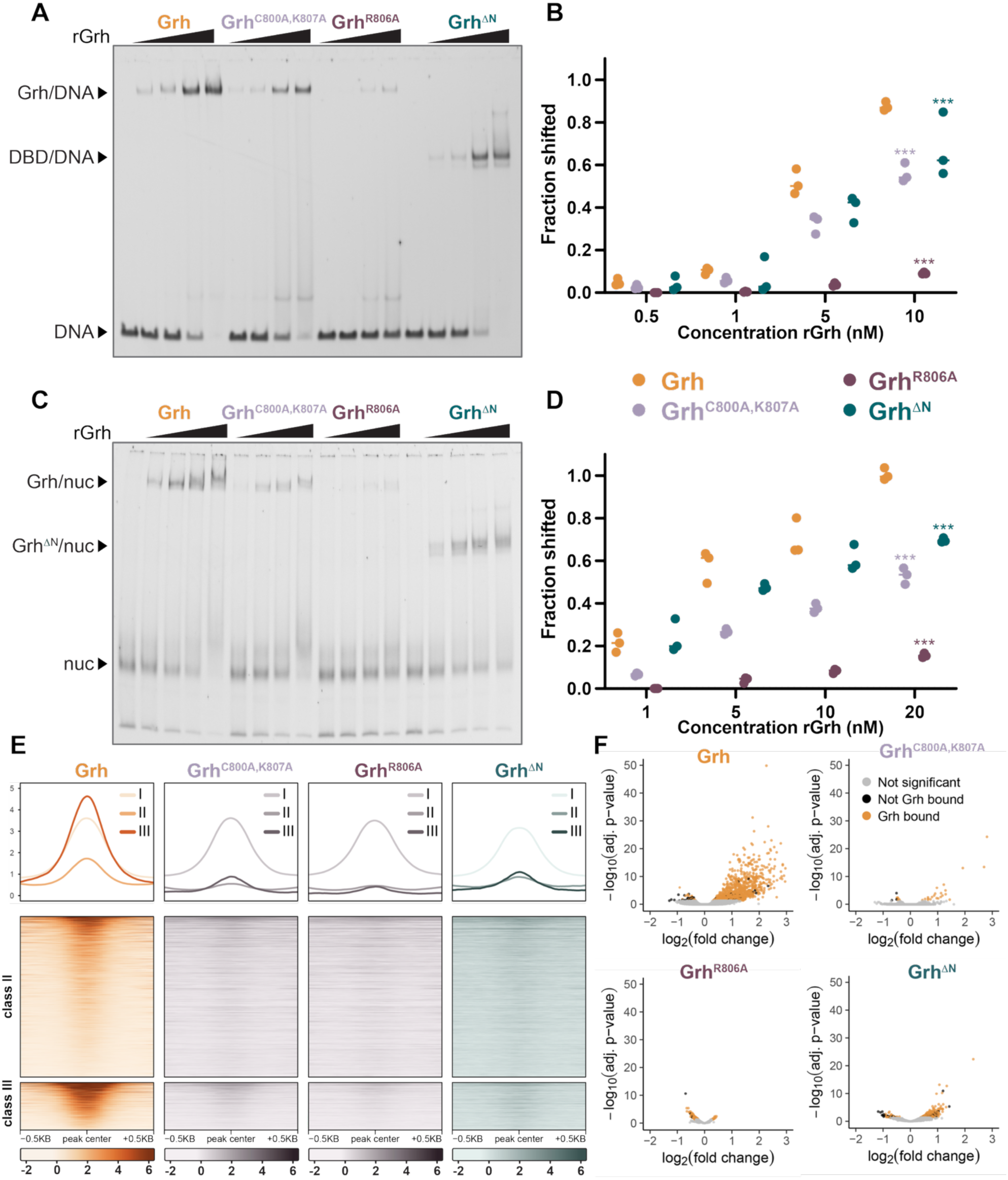
Grainy head requires the DNA-binding domain and regions outside for binding nucleosomes and opening chromatin. (**A**) EMSA with increasing concentrations of recombinant Grh (rGrh), Grh^C800A,K807A^, Grh^R806A^ or Grh^ΔN^ (0 – 10 nM) incubated with Cy5-labeled DNA. (**B**) Quantification of the fraction of Grh-bound DNA, indicated by the proportion shifted in the EMSA shown in (A) and two additional experiments. Significance compared to rGrh was determined with a two-way ANOVA with post-hoc Dunnett’s multiple comparisons test and, for simplicity, is show for the highest concentration tested. ***, p<0.0005 (**C**) EMSA with increasing concentrations of rGrh, Grh^C800A,K807A^, Grh^R806A^ or Grh^ΔN^ (0 - 20nM) incubated with Cy5-labeled nucleosomal DNA. (**D**) Quantification of the fraction of Grh-bound nucleosomes, indicated by the proportion shifted in the EMSA shown in (C) and two additional experiments. Significance compared to rGrh was determined with a two-way ANOVA with post-hoc Dunnett’s multiple comparisons test and, for simplicity, is shown only for the highest concentration tested. ***, p<0.0005 (**E**) Heatmaps and metaplots of ChIP-seq for Grh, Grh^C800A,K807A^, Grh^R806A^ or Grh^ΔN^ induced in S2 cells. Class I shown in metaplots, Class II and III shown in both metaplots and heatmaps. (Grh^ΔN^ data from Gibson et al. 2024 (*12*)). (**F**) Volcano plots of changes in ATAC-seq signal upon expression of Grh, Grh^C800A,K807A^, Grh^R806A^ or Grh^ΔN^ as compared to uninduced controls. (Grh^ΔN^ data from Gibson et al. 2024 (*12*)). Regions bound by wild-type Grh, as defined in Gibson et al. 2024 (*12*), shown in orange.

To determine how binding to nucleosomes *in vitro* correlated with pioneering activity in cells, we expressed both DBD mutants and Grh^ΔN^ in S2 cells and assayed binding and chromatin accessibility. All three proteins retained binding to most class I sites, reflecting their ability to occupy accessible chromatin (Fig. 2E, Fig. S3B) (*12*). Both Grh^C800A,K807A^ and Grh^R806A^ had reduced binding to accessible sites containing the Grh motif, supporting the role of these residues in promoting sequence-specific Grh binding (Fig. 2E, Fig. S3B). In contrast to their ability to bind to open chromatin, all three proteins had markedly decreased binding to closed chromatin (both class II and class III sites) (Fig. 2E) (*12*). Consistent with the loss of binding at inaccessible sites, these mutants failed to open chromatin, despite being expressed at levels higher than wild type (Fig. 2F, Fig. S3C, D) (*12*). Thus, while Grh^C800A,K807A^ and Grh^ΔN^ had modest effects on binding *in vitro*, the reduction in chromatin binding was much more pronounced when assayed in cells. We next evaluated the contributions of these domains to transcriptional activation using a luciferase reporter driven by the promoter for *homogentisate 1,2 dioxygenase* (*hgo*), an established Grh target containing the Grh motif (*68*). Both DBD mutants and Grh^ΔN^ failed to activate the reporter, indicating that the structured DBD and the disordered N-terminus are required to promote transcription (Fig. S3E, F). By assaying the same constructs both *in vitro* and in cells, we demonstrate that mononucleosome binding only partially captures the requirements for pioneering and that pioneering activity requires both DNA binding and activities provided by the N-terminus.

### Grh requires the DBD and regions outside for stable chromatin occupancy

We hypothesized that the disconnect between mononucleosome binding and pioneering activity in cells might arise from differences in protein dynamics *in vivo*. Both EMSAs and ChIP-seq provide static views of protein binding. We therefore sought to use *in vivo* imaging to determine how the DBD and N-terminus contribute to the dynamics of Grh binding. Because Grh binds DNA as a dimer, we expressed the mutants in a tissue that lacks endogenous protein to ensure we were not assaying binding of mixed dimers (*30*, *47*). For this reason, we chose to express Halo-tagged Grh constructs in *Drosophila* salivary glands, which have large polytene chromosomes that can be easily visualized in intact nuclei and are ideal for live imaging of chromatin dynamics (*69*). To assay proteins that show a range of binding in our prior assays (Fig. 2), we expressed Halo-tagged wild-type Grh, Grh^R806A^, or Grh^ΔN^ and analyzed chromosomal occupancy following incubation with the Halo ligand JF646. Wild-type Grh was strongly chromosome-associated and was localized to multipole distinct bands, corresponding to individual genomic loci (Fig. 3A). Grh^ΔN^ was enriched on the chromosomes but with less clear localization to distinct bands than the full-length protein and with some protein remaining diffuse in the nucleoplasm (Fig. 3A). By contrast, Grh^R806A^ was present in the nucleoplasm but appeared to be largely excluded from chromosomes (Fig. 3A). We then performed fluorescent recovery after photobleaching (FRAP) to quantitatively assess the mobility of Grh on chromatin. We assayed wild-type Grh and Grh^ΔN^, focusing on the chromatin-bound fraction that was visualized through colocalization with the chromosomes. Grh^R806A^ was excluded from analysis as it was not detectably present on chromosomes. Wild-type Grh recovered slowly (t^1/2^ = 291s), indicating a slow exchange of molecules, and never achieved full recovery within the imaging window, plateauing at 50%. This indicates that there is a substantial fraction of the protein that is essentially immobile over the time-course of the experiment (Fig. 3B). Grh^ΔN^ recovered dramatically faster (t^1/2^ = 21s) and reached close to 100% recovery, indicating that the majority of chromosomally associated mutant proteins are rapidly exchanging. Hence, the N-terminus is required for stable chromatin binding (Fig. 3B).

**Figure 3.**
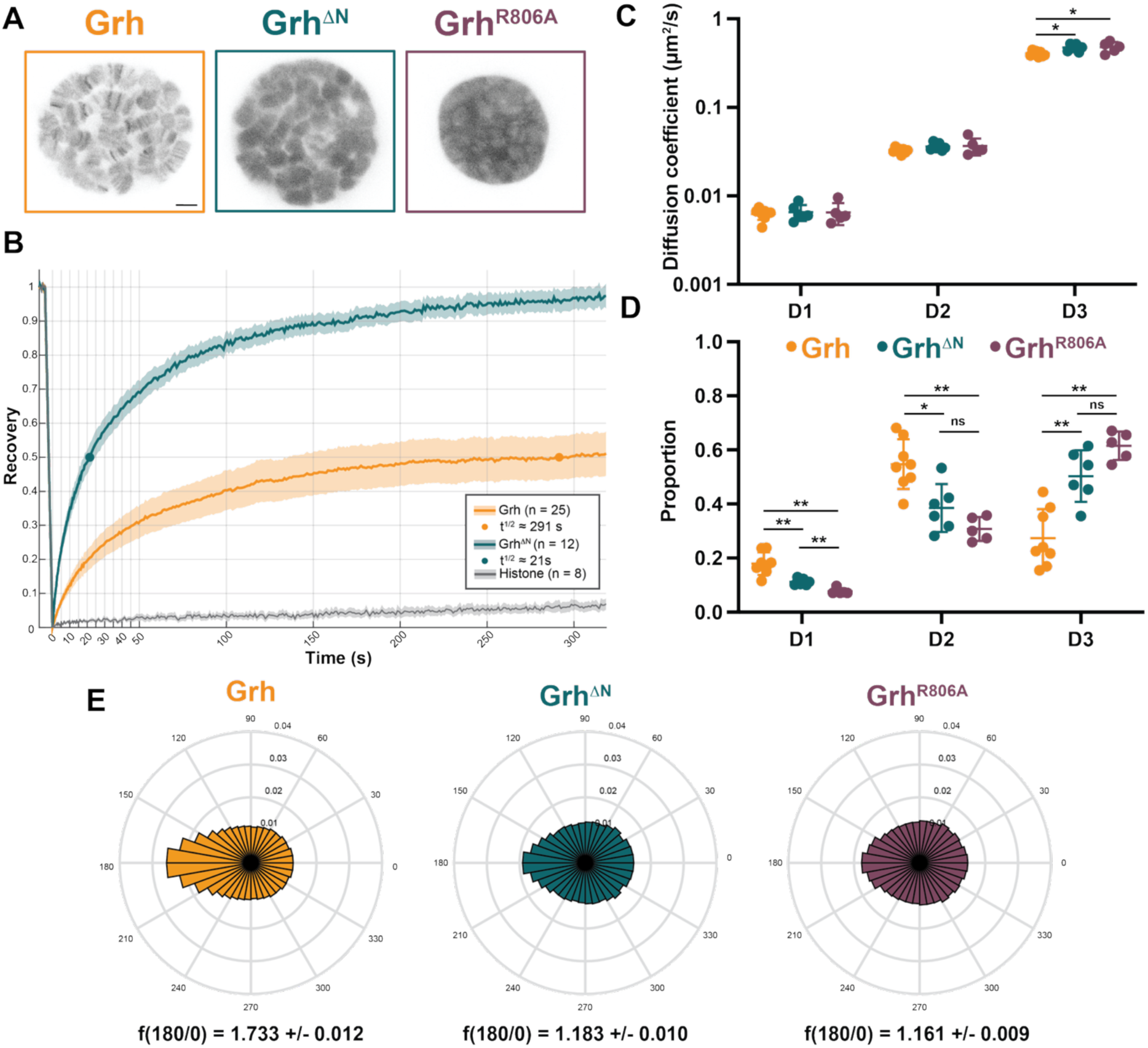
Grainy head requires the DNA-binding domain and regions outside for stable chromatin occupancy. (**A**) Representative images of single nuclei of third instar larval salivary glands labelled with J646 to visualize expression of Halo-tagged Grh, Grh^ΔN^ or Grh^R806A^. Scale bar, 5 μm. All images at the same magnification. (**B**) Recovery of Grh or Grh^ΔN^ molecules after photobleaching. Legend summarizes numbers of nuclei (n) and time to 50% recovery (t^1/2^). Shading indicates the standard error of the mean (SEM). (**C**) Average diffusion coefficients of three Grh populations as determined by vbSPT. Significance was assessed with two-sample *t* tests. *, p < 0.05 Unmarked comparisons showed no statistical difference. (**D**). Average proportion of Grh, Grh^R806A^ or Grh^ΔN^ molecules per nucleus found in each of the three states determined by vbSPT. (Grh, n = 8 nuclei, 103,847 trajectories; Grh^R806A^, n = 5 nuclei, 39,946 trajectories; Grh^ΔN^, n = 6 nuclei, 50,109 trajectories). Significance was assessed with Mann-Whitney *U* tests. *, p < 0.05 **, p < 0.005 (**E**) Circular histograms of angle distributions calculated from D3 trajectories. Fold anisotropy f(180/0) ± SD is noted for each sample.

To further investigate the dynamics of Grh binding, we performed single-particle tracking (SPT) using sparse labelling of the Halo-tagged proteins. Molecules were imaged at 50 ms per frame, and their trajectories were tracked and subsequently analyzed using a three-state Hidden Markov model, which assumes that each Grh molecule exists in one of three distinct states, each with a specific diffusion coefficient. The resulting states were: D1, the slowest diffusing population likely corresponding to stably bound molecules; D2, an intermediate population whose diffusion coefficient is consistent with less stably bound molecules; D3, the most rapidly diffusing population, that is likely composed of freely diffusing and transiently interacting molecules. Diffusion coefficients for each state were similar between wild-type Grh, Grh^R806A^ and Grh^ΔN^ populations, confirming that the same population characteristics were sampled in all conditions (Fig. 3C). In comparison to wild-type Grh, the proportions of Grh^R806A^ and Grh^ΔN^ molecules in the D1 and D2 state were considerably reduced, consistent with less stable chromatin engagement (Fig. 3D). In agreement with the lack of enrichment on chromosomes, very few Grh^R806A^ molecules exhibited D1 state properties (Fig. 3D). To further examine how the DBD and N-terminus contribute to Grh chromatin recruitment, we measured the angles within the trajectories of the most dynamic molecules (D3). Any deviation of the angle distributions from uniform is indicative of behaviors that differ from Brownian motion, such that molecules with the more confined behaviors indicative of searching behavior have greater angular anisotropy. Indeed wild-type Grh molecules had a high degree of angular anisotropy while both Grh^ΔN^ and Grh^R806A^ displayed reduced anisotropy, demonstrating less constrained movement and a decrease in searching behavior (Fig. 3E). Our quantitative assessment of how the DBD and N-terminus contribute to the dynamics of Grh chromatin occupancy shows that DNA binding is essential for chromatin occupancy. By contrast, the disordered N-terminus of Grh supports stable binding and aids local searching but is not essential for chromatin association.

### The intrinsically disordered N-terminus contributes to pioneering independent of amino acid sequence

Given the critical role of the DBD in mediating DNA contacts, its necessity for pioneering activity was expected. By contrast, the mechanisms by which the disordered N-terminus contributes to stable chromatin occupancy and pioneering activity were less clear. The N-terminus is highly disordered apart from a short-structured domain (SD) that has previously been implicated in transcriptional activation (Fig. 4A) (*47*). To determine whether the SD was required for transcriptional activation in our assays, we tested whether Grh lacking this domain (Grh^ΔSD^) could promote expression from the *hgo* promoter. In contrast to expectations, deletion of the SD did not impact Grh-mediated transcriptional activation (Fig. 4B, Fig. S4B). Thus, we largely focused on the disordered region of Grh. Because intrinsically disordered domains are a common feature among eukaryotic transcription factors and have been associated with pioneering activity, we asked whether the importance of the N-terminal domain reflects specific-sequence elements (*15*, *17*, *20*, *70*). We generated a scrambled N-terminal mutant (Grh^Scr^) that preserves the overall amino acid composition and predicted disorder but disrupts the sequence, while leaving the C-terminal DBD and dimerization domain intact (Fig. 4A). Scrambling the N-terminal sequence did not substantially alter the distribution of charge or hydropathy (Fig. S4A). For consistency, we selected a scrambled sequence that retained a patch of predicted order around the same size and location as the SD in the wild-type protein but with an entirely scrambled amino acid sequence.

**Figure 4.**
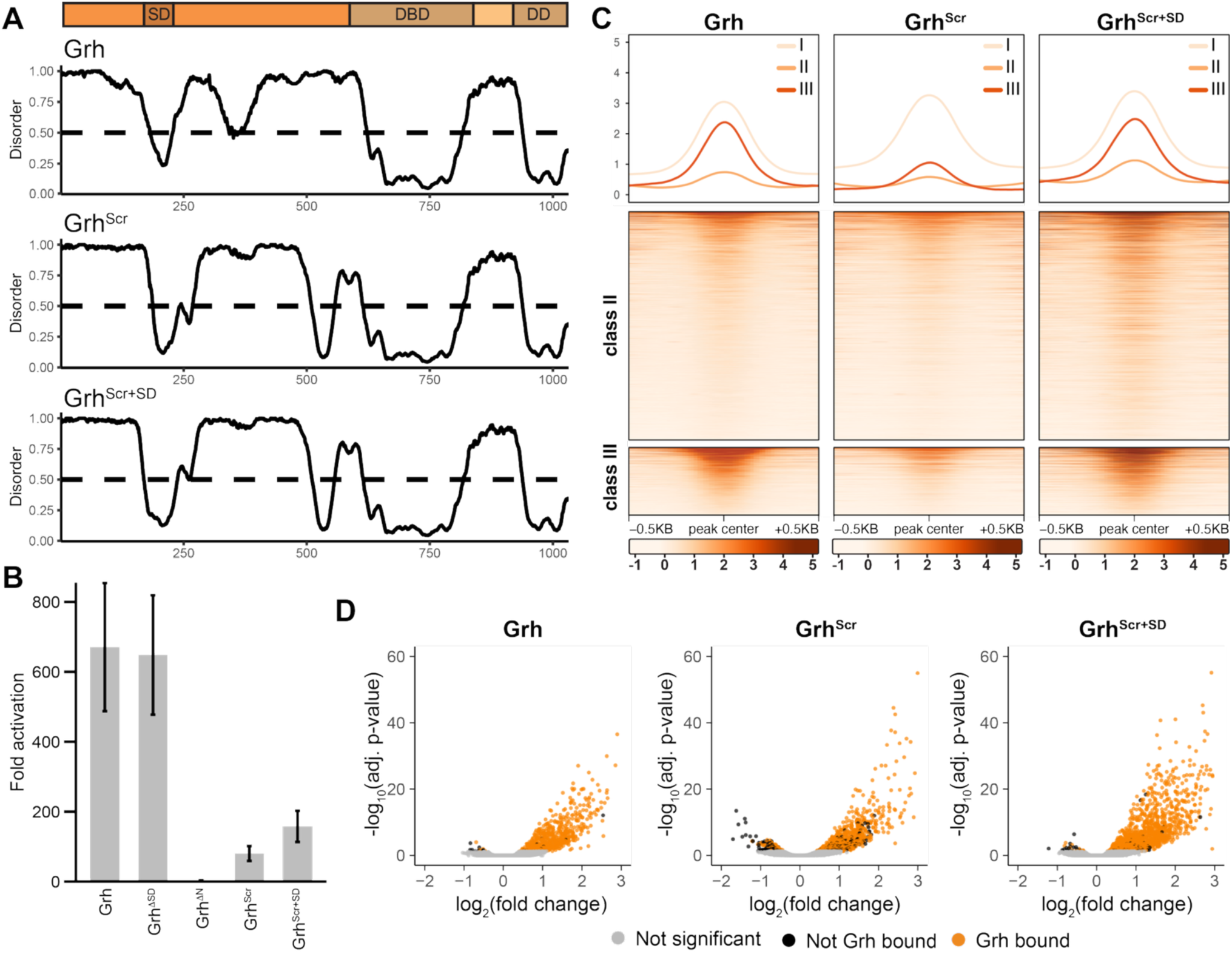
The intrinsically disordered Grh N-terminus can contribute to pioneering independent of amino acid sequence. (**A**) Graph of predicted disorder score (from Metapredict (*86*)), Grh (top), Grh^Scr^ (center), Grh^Scr+SD^ (bottom). (**B**) Fold activation of the *hgo* luciferase reporter by either Grh, Grh^ΔSD^, Grh^ΔN^, Grh^Scr^, or Grh^Scr+SD^. n=3, mean ± S.D. (**C**) Heatmaps and metaplots of ChIP-seq for Grh, Grh^Scr^, or Grh^Scr+SD^ expressed in S2 cells. Class I shown in metaplots, Class II and III shown in both metaplots and heatmaps. (**D**) Volcano plots of changes in ATAC-seq signal upon expression of Grh, Grh^Scr^, or Grh^Scr+SD^ as compared to uninduced controls. Regions bound by wild-type Grh, as defined in Gibson et al. 2024 (*12*), shown in orange.

We tested the ability of Grh^Scr^ to activate transcription using the Grh-responsive *hgo* reporter. Although the levels of Grh^Scr^ were consistently lower than those of wild-type Grh, the scrambled N-terminus partially restored transcriptional activation activity relative to Grh^ΔN^ (Fig. 4B, Fig. S4B). Because the ability to activate transcription does not reflect pioneering activity, we directly assessed chromatin binding and opening using ChIP-seq and ATAC-seq, respectively. Grh^Scr^ exhibited reduced binding at inaccessible sites, similar to Grh^ΔN^ (Fig. 2E, Fig. 4C) (*12*). However, whereas Grh^ΔN^ failed to open chromatin, the addition of the scrambled N-terminus allowed Grh^Scr^ to open chromatin, albeit not to the degree of wild-type Grh (Fig. 2F, Fig. 4D) (*12*). Although Grh^Scr^ was expressed at slightly higher levels than wild type, Grh^ΔN^ fails to open chromatin even when overexpressed, indicating that the disordered N-terminus in Grh^Scr^ is sufficient to partially restore pioneering activity (Fig. S4C) (*12*). Reintroduction of the structured domain into the scrambled N-terminus to generate Grh^Scr+SD^ improved the capacity of Grh to activate transcription, bind closed chromatin and promote chromatin accessibility (Fig. 4B-D). Thus, the SD sequence, rather than structure alone, promotes pioneer activity. However, the capacity of the scrambled N-terminus to confer partial activity suggests that sequence-specific features are not strictly required, and other properties of the disordered region support some aspects of pioneering.

### Diverse N-terminal domains are sufficient for pioneering

The requirement of the N-terminal domain for pioneering in *Drosophila* Grh contrasts with the widespread lack of conservation in this domain across species (*48*). While the mammalian GRHL family of proteins share the deeply conserved C-terminal DNA-binding and dimerization domain, the N-terminal domain is truncated and less disordered than *Drosophila* Grh (Fig. S5A) (*33–35*). To expand on the properties of the N-terminus that contribute to pioneering we asked whether human GRHL2 could function in *Drosophila* cells. Using the *hgo* promoter, we demonstrated that GRHL2 failed to activate transcription (Fig. 5A, Fig. S5B). This lack of activity could reflect either a failure of GRHL2 to bind DNA in *Drosophila* cells or a lack of recruitment of cofactors needed to promote transcription. We therefore tested the pioneering capacity of GRHL2 using ChIP-seq and ATAC-seq on S2 cells expressing human GRHL2. GRHL2 was N-terminally tagged with an HA epitope since the Grh antibody does not recognize human GRHL2. As a control, we similarly tagged *Drosophila* Grh with HA. 99.6% of ChIP peaks identified with the HA antibody overlapped with peaks identified using antibodies recognizing Grh, demonstrating the specificity of the pulldowns (Fig. S5C). In contrast to the failure of GRHL2 to promote gene expression from the *hgo* promoter, GRHL2 bound to 9,533 loci, including substantial binding to both class II and class III regions of closed chromatin occupied by Grh (Fig. 5B, Fig. S5D). By comparing binding between Grh and GRHL2 we found that the shared sites had the highest ChIP signal for each protein and were enriched for the canonical Grh-binding motif (Fig. S5E, F). Sites bound uniquely by GRHL2 contained a more degenerate motif (Fig. S5F). GRHL2 not only binds inaccessible sites, but ATAC-seq revealed that GRHL2 induces chromatin accessibility at these sites, although not to the extent of *Drosophila* Grh (Fig. 5C). Thus, human GRHL2 can function as a pioneer factor in *Drosophila* cells, but this activity is not sufficient to promote gene expression.

**Figure 5.**
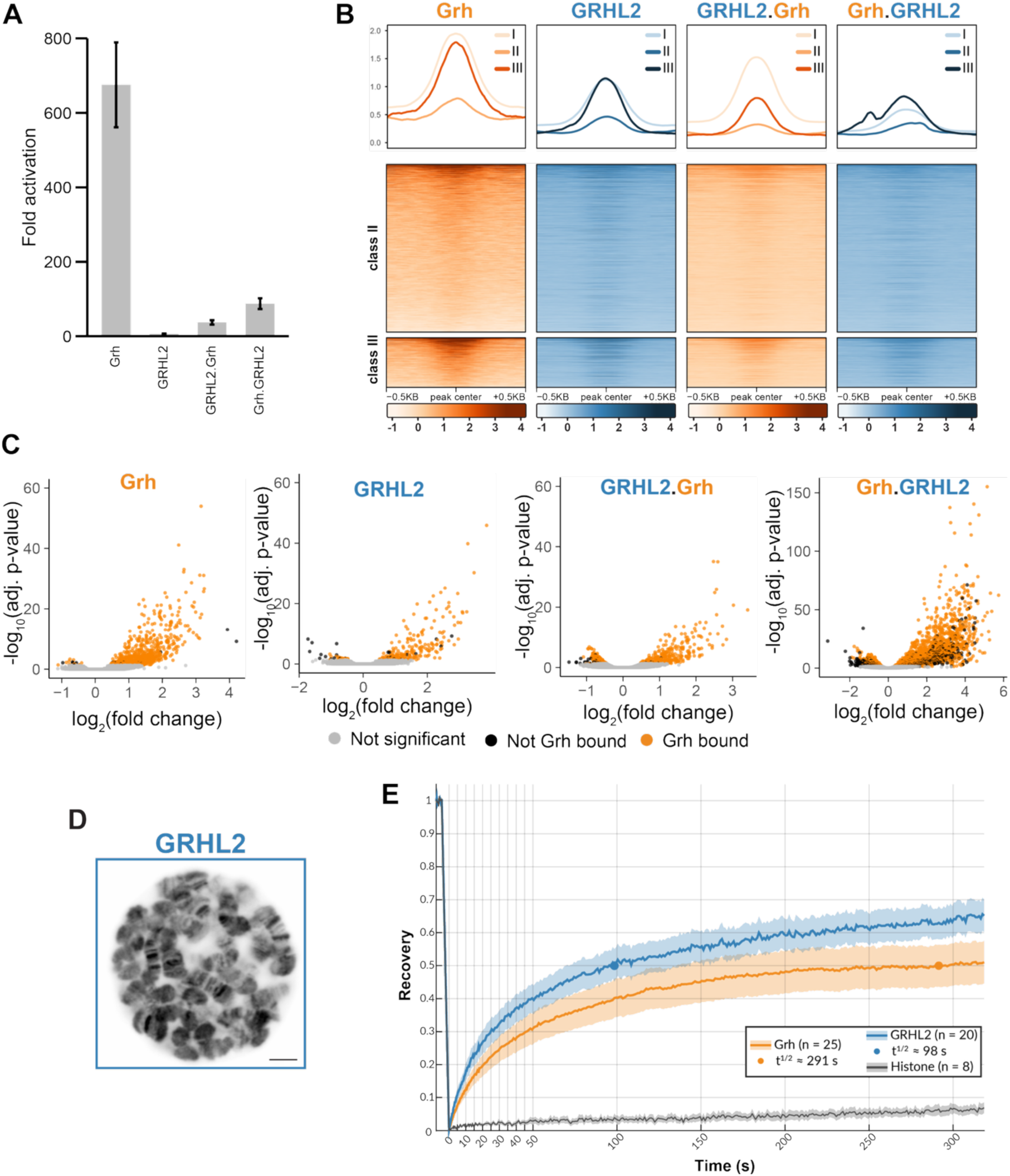
Diverse N-terminal domains are sufficient for pioneering. (**A**) Fold activation of the *hgo* luciferase reporter by either *Drosophila* Grh, human GRHL2 or chimeras (GRHL2.Grh, Grh.GRHL2). (**B**) Heatmaps and metaplots of ChIP-seq for Grh, GRHL2, or chimeras expressed in S2 cells. Class I shown in metaplots, Class II and III shown in both metaplots and heatmaps. (**C**). Volcano plots of changes in ATAC-seq signal upon expression of *Drosophila* Grh, human GRHL2 or chimeras as compared to uninduced controls. Regions bound by wild-type Grh, as defined in Gibson et al. 2024 (*12*), shown in orange. (**D**) Representative images of single nuclei of third instar larval salivary glands labelled with J646 to visualize expression of Halo-tagged GRHL2. Scale bar, 5 μm. (**E**) Recovery of Grh or GRHL2 molecules after photobleaching. Legend summarizes numbers of nuclei (n) and time to 50% recovery (t^1/2^). Shading indicates the standard error of the mean (SEM).

Our quantitative imaging data suggested that the disordered N-terminus stabilized chromatin occupancy and that this was correlated with pioneering activity. To determine if the pioneering activity of human GRHL2 was similarly associated with stable chromatin occupancy, we expressed Halo-tagged GRHL2 in salivary gland nuclei. Supporting the ChIP-seq results that GRHL2 binds chromatin in *Drosophila*, we saw robust localization to the polytene chromosomes (Fig. 5D). FRAP demonstrated that human GRHL2 recovered with kinetics similar to *Drosophila* Grh with a substantial immobile fraction, reflecting stable chromatin binding (Fig. 5E). Together the imaging and genomics analyses shows that despite possessing a substantially shorter and more ordered N-terminus than *Drosophila* Grh, human GRHL2 can stably bind closed chromatin and promote accessibility when expressed in *Drosophila* cells.

To further explore the individual contributions of the C-terminal versus N-terminus, we generated chimeric fusions between the human and *Drosophila* Grh proteins: GRHL2.Grh, in which the N-terminal domain of GRHL2 is fused to the *Drosophila* Grh DNA-binding and dimerization domains, and Grh.GRHL2, in which the *Drosophila* Grh N-terminal domain is fused to the GRHL2 DNA-binding and dimerization domains. Both chimeric proteins weakly activate the *hgo* reporter, but not nearly to the extent of *Drosophila* Grh (Fig. 5A, Fig. S5B). ChIP-seq and ATAC-seq demonstrated that that the GRHL2.Grh chimera bound closed chromatin and promoted accessibility to a similar extent as GRHL2 (Fig. 5B,C). The Grh.GRHL2 fusion also bound closed chromatin to a similar extent as GRHL2 but promoted accessibility to a much greater degree (Fig. 5B,C). Quantitative comparisons to *Drosophila* Grh were complicated by the relatively higher expression levels of GRHL2 and the fusion proteins (Fig. S5D). Overall, these results indicate that while the human GRHL2 N-terminus can confer pioneering activity to Grh, the extended *Drosophila* N-terminal is most efficient at promoting chromatin accessibility in *Drosophila* cells.

### Mitotic retention of Grh requires DNA binding

We further sought to connect pioneer activity with mitotic retention by testing the role of both the DBD and extended N-terminus in retention of Grh on mitotic chromosomes. We examined mitotic retention of GFP-tagged Grh, Grh^ΔN^, Grh^R806A^ and GRHL2 in S2 cells, marking mitotic chromosomes with mCherry-tubulin. Grh^R806A^ was not mitotically retained, consistent with the importance of direct DNA contacts for pioneering and demonstrating that these contacts are also required for mitotic retention (Fig. 6). This lack of retention is not due to the change in charge, as substitution of the arginine with either lysine or histidine similarly abrogated mitotic retention (Fig. S6). Grh^ΔN^ showed robust retention, in stark contrast to the requirement of the N-terminus for stable chromatin interactions and efficient pioneering activity (Fig. 6). Indeed, GRHL2 was similarly retained (Fig. 6). Together these results demonstrate that mitotic retention can be uncoupled from the stable chromatin occupancy required for pioneering activity but depends on DNA binding.

**Figure 6.**
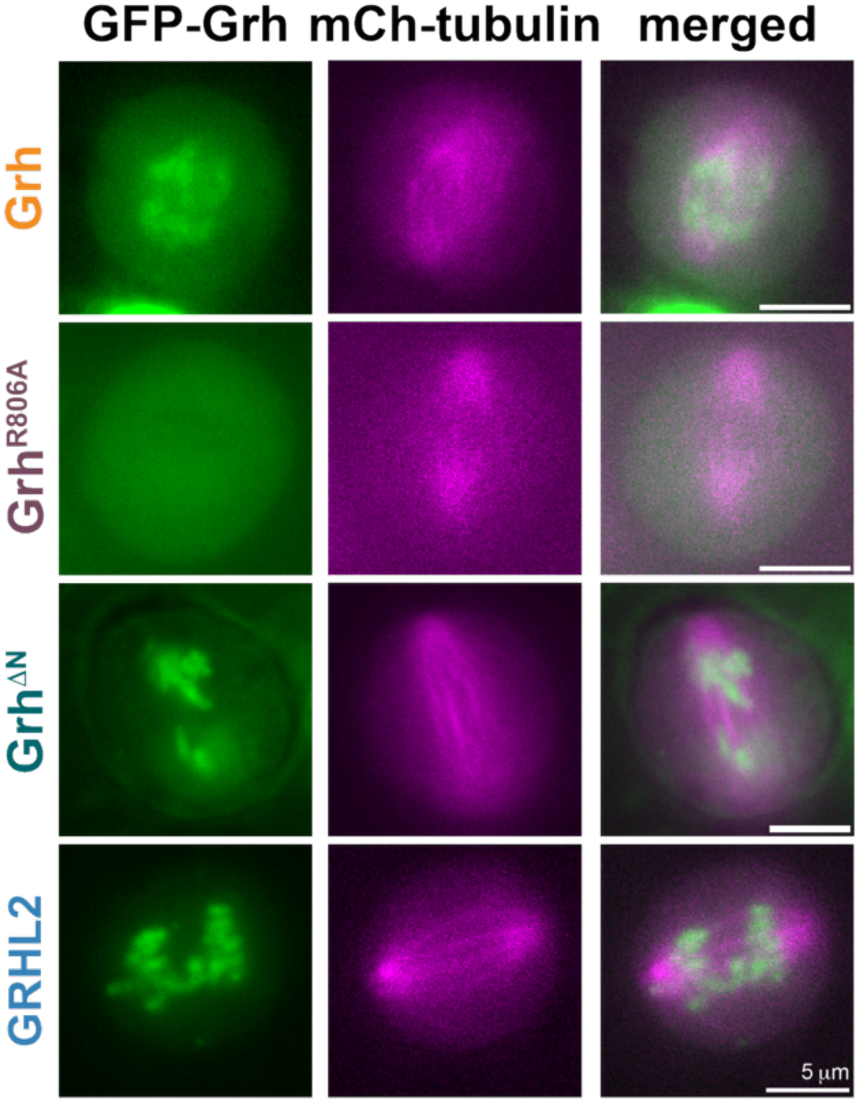
Mitotic retention of Grainy head requires only the C-terminal domains. Images of GFP-tagged Grh proteins (green) and mCherry-tubulin (magenta) at anaphase in S2 cells.

## DISCUSSION

Pioneer transcription factors are defined by their ability to engage nucleosomal DNA and initiate chromatin opening, enabling them to act at the top of gene-regulatory networks. Nonetheless, the essential defining features and how they contribute to the unique properties of pioneer factors remain controversial. Proteins with pioneering characteristics do not share DBD structures or sequences outside the DBD, adding further complexity to the challenge of defining this specialized class of transcription factors. To begin to address this complexity, we used multiple defining assays of pioneering function to identify the properties of the deeply conserved pioneer factor Grh. By doing so, we determined how these features relate to each other and how they rely on specific protein domains. Our assays reveal distinct requirements for pioneering and mitotic retention. Sequence-specific DNA binding, but not the multivalent interactions mediated by the disordered N-terminus, is required for retention of Grh on mitotic chromatin. Thus, the set of intrinsic features enabling stable chromatin engagement during interphase is larger than that supporting chromatin associations during mitosis. This distinction is further supported by the fact that mitotic retention is not shared by a subset of pioneer factors, including Zelda and Ascl1 (*71*, *72*).

The DNA-binding domain of Grh defines a family of transcription factors conserved from fungi to humans that bind to a shared DNA-sequence motif. In addition, DNA binding by Grh depends on dimerization mediated by a conserved dimerization domain (*30*, *47*). DBDs are classically thought of as the primary drivers of nucleosome targeting, and *in vitro*, are often sufficient for nucleosome binding (*7*, *9*, *10*). While Grh-family members were previously shown to bind and open closed chromatin, the ability to engage nucleosomes *in vitro* had not been investigated (*12*, *44–46*). Like many pioneer factors, Grh binds nucleosomes at the entry/exit site with affinities similar to that of free DNA. This interaction is largely mediated through sequence-specific binding by the C-terminal DBD and dimerization domains. However, the extended, disordered N-terminus also contributes. This contrasts with the requirement for the N-terminus for chromatin-binding in cell culture, demonstrating that *in vitro* binding to mononucleosomes only partially reflects the requirements of pioneer factors to occupy the more complex chromatin environment in cells and is consistent with similar results for other pioneer factors (*13*). By combining these static assays of pioneer function with investigation of how protein domains contribute to protein dynamics *in vivo*, our data support an essential role for the N-terminus in promoting the stable chromatin occupancy necessary for binding closed, but not accessible chromatin. Furthermore, both domains promote localized chromatin scanning, indicated by less compact trajectories when these domains are mutated. Together, these data demonstrate that the structured DBD and the disordered N-terminus are required in combination for the stable chromatin occupancy required for pioneer function. These requirements likely reflect complex engagements with chromatin that are only partially reflected in the ability to bind mononucleosomes *in vitro*.

Despite contributing to Grh pioneering function, the N-terminus is largely unstructured and lacks sequence conservation. There is a single, structured domain whose function is unclear. While it was initially defined as the minimal transactivation domain in *Drosophila* cells, when expressed in yeast this domain does not activate reporter gene expression (*47*). Furthermore, we showed that the minimal transactivation domain (SD) is not required for activation of a reporter gene nor is it sufficient to promote wild-type levels of activation. While these data demonstrate that the structured domain is not, as previously suggested, the only transactivation domain, addition of this domain to a scrambled N-terminus increased binding and opening of closed chromatin, suggesting it promotes pioneer activity.

Apart from the structured region, the N-terminus contributes to pioneering through a mechanism that is, at least in part, independent of amino acid sequence. This activity is similar to what has been demonstrated for activation domains of yeast transcription factors, two-thirds of which retain the ability to activate gene expression when scrambled (*73*). Disordered regions are a shared feature of many eukaryotic transcription factors and contribute to chromatin binding through multiple mechanisms including directly engaging with chromatin and facilitating increased local concentrations through multivalent interactions. While the N-terminus of human GRHL2 is shorter and has less predicted disorder than *Drosophila* Grh, it enables the stable chromatin occupancy required for binding and opening closed chromatin. Similarly, the *C. elegans* Grh ortholog, which is also truncated relative to *Drosophila* Grh, partially rescues embryonic viability and chromatin accessibility in a *Drosophila grh*-null background (*48*). These data support a role for weak, non-specific, multivalent interactions mediated by disordered N-termini in promoting pioneer activity. By contrast, the scrambled N-terminus of Grh and the N-terminus of GRHL2 do not allow for activation of a reporter gene, separating pioneering from transcriptional activation and suggesting that sequence-specific protein interactions may be required for activation.

Using multiple, complementary assays, we define how protein-intrinsic features contribute to the defining features of the pioneer factor Grh. Both sequence-specific interactions with DNA, driven by the conserved DBD, and non-specific interactions, mediated by the unstructured N-terminus, contribute to the stable chromatin occupancy required for Grh pioneer-factor activity. We suggest that despite having distinct polypeptide sequences, pioneer factors require stable chromatin occupancy, which can be mediated through diverse mechanisms. Many pioneer factors, including Grh, are dysregulated in cancers and discrete protein domains can be brought together through oncogenic fusions. Thus, our analysis of how multiple protein domains contribute to pioneer activity has important implications for both normal development and cancer.

## MATERIALS AND METHODS

### Experimental design

#### Nucleosome reconstitution

The 159bp endogenous DNA fragment (dm3_chr2R: 19,981,929-19,982,087) was amplified from *Drosophila* genomic DNA. For Widom 601 nucleosomes, the Grh motif (AACCGGTT) was inserted into the 153bp Widom 601 sequence at super helical locations (SHL) -6.5, -4.5, -4.0, and 0 (the dyad) (*51*). DNA was amplified using one Cy5-labeled primer and one unlabeled primer. Widom 601 without the Grh motif was used as a control and amplified using one Cy3-labeled primer and one unlabeled primer. The DNA was ethanol precipitated and purified with AxyPrep Mag PCR Clean-up beads (Axygen) using a 1.8x ratio of beads to sample. The nucleosomes were reconstituted using salt gradient dialysis (*74*). The Grh motif is underlined, and mutated base pairs (from canonical Widom 601 sequence) are shown in lowercase.

Endogenous DNA:

Cy5-CCATTTTGGCGGTGGATCCAAGCGGCCAAGTGCTAATGCCCCCAGCCCCGCCGATTGCTATGTCTTT GAGCTCCAACTTGTCACGCTGCCACAGACTGGAGCTCTCTCTCCGCGAA<u>AACCGGTT</u>CTCTGGTGGCCG GTCGCAAACGAAATCTCCATCTG

Widom 601 DNA:

Cy3-ATCCTGGAGAATCCCGGTGCCGAGGCCGCTCAATTGGTCGTAGACAGCTCTAGCACCGCTTAAACGCACGTACGCGCTGTCCCCCGCGTTTTAACCGCCAAGGGGATTACTCCCTAGTCTCCAGGCACGTGTCAGATATATACATCCTGTGAT

Widom 601 DNA with Grh motif at SHL-6.5

ATCCTGaAccggttCGGTGCCGAGGCCGCTCAATTGGTCGTAGACAGCTCTAGCACCGCTTAAACGCACGTACGCGCTGTCCCCCGCGTTTTAACCGCCAAGGGGATTACTCCCTAGTCTCCAGGCACGTGTCAGATATATACATCCTGTGAT-Cy5

Widom 601 DNA with Grh motif at SHL-4.5

ATCCTGGAGAATCCCGGTGCCGAGGCCaacCggTTGGTCGTAGACAGCTCTAGCACCGCTTAAACGCACGTACGCGCTGTCCCCCGCGTTTTAACCGCCAAGGGGATTACTCCCTAGTCTCCAGGCACGTGTCAGATATATACATCCTGTGAT-Cy5

Widom 601 with Grh motif at SHL-4.0

ATCCTGGAGAATCCCGGTGCCGAGGCCGCTCAAccGGtTGTAGACAGCTCTAGCACCGCTTAAACGCACGTACGCGCTGTCCCCCGCGTTTTAACCGCCAAGGGGATTACTCCCTAGTCTCCAGGCACGTGTCAGATATATACATCCTGTGAT-Cy5

Widom 601 with Grh motif at SHL0

ATCCTGGAGAATCCCGGTGCCGAGGCCGCTCAATTGGTCGTAGACAGCTCTAGCACCGCTTAAACGCACG T<u>AacCGgTt</u>TCCCCCGCGTTTTAACCGCCAAGGGGATTACTCCCTAGTCTCCAGGCACGTGTCAGATATATAC ATCCTGTGAT-Cy5

#### Protein expression and purification

As described previously, MBP-tagged Grh^ΔN^ (Grh^603-1032^) was purified from *E. coli* (*75*, *76*) with the exclusion of the final dialysis step. Briefly, protein was bound to amylose resin (New England Biolabs) and eluted with 20 mM maltose. Baculovirus expression was used to purify full-length Grh-PH as described previously (*75*). For purification of Grh^C800A, K807A^ and Grh^R806A^ cDNA encoding each mutant was cloned into pDEST8 and used to generate baculovirus using the Bac-to-Bac expression system (ThermoFisher). Freshly amplified virus was used to infect Sf9 cells grown in cytiva SFX media (Fisher Scientific) supplemented with 10% fetal bovine serum. As in McDaniel et al. 2019 (*11*), three days following infection cells were collected on ice, centrifuged, washed with PBS containing 5 mM MgCl2, and centrifuged once more. The cells were resuspended in hypotonic buffer (15 mM HEPES, 15 mM KCl, 2 mM MgCl2, 0.02% Tween, 10% glycerol, 2 mM BME, 1 mM EGTA, 0.4 mM PMSF, Pierce protease inhibitor tablets (Fisher)), flash frozen in liquid nitrogen, and kept at -80°C. Cell suspensions were thawed and dounced. KCl was then added to bring the concentration to 300 mM and the lysate was cleared by centrifugation (10,000 rpm for 10 min at 4°C, 16,000rpm for 10 min at 4°C). 20 µM PMSF and 750 µL pre-washed anti-flag M2 affinity beads (Sigma A2220) were added to the extract and incubated at 4°C for 3 hours. The beads were washed, and protein was eluted in buffer containing 150 mM KCl and 200 µg/mL Flag peptide following a 30 min incubation with end-over end mixing at 4°C. Protein concentration was determined by comparing levels to BSA standards using Coomassie blue staining.

#### Electrophoretic Mobility Shift Assays

2.5 nM (50 fmol/ 20 µl reaction) of Cy5-labelled probe (DNA or nucleosome) was incubated with recombinant Grh protein in buffer containing: 5 ng poly[d-(IC)], 12.5 mM HEPES, 0.5 mM EDTA, 0.5 mM EGTA, 5% glycerol, 0.25 mM DTT, 150 μM PMSF, 0.075 mg/ml BSA, 2.5 mM MgCl2, 0.005% Tween, 50 mM KCl at room temperature for 60 min (30 min for Fig. 1A). For reactions containing both Cy5- and Cy3-labeled Widom 601 DNA or nucleosomes, each was present at 2.5 nM (5 nM total). Reactions were run on a 4% non-denaturing polyacrylamide gel in 0.5x Tris Borate EDTA. Gels were visualized with a Typhoon FLA9000 using Cy5 and Cy3 fluorescence settings. The fraction of bound protein was quantified by dividing the shifted band of each respective reaction by the unshifted band from the no protein reaction. Quantification was performed on 3 or 4 independent gels.

#### Cell culture and generation of stable cell lines

We used a stable cell line expressing full-length wild type Grh that was generated previously (*12*). There is an alternative ATG 252 bp after the initial start codon, resulting in multiple protein products upon copper induction.

Additional expression constructs were generated as described below and cloned into pMT-puro. Point mutants, Grh^C800A, K807A^ and Grh^R806A^ were generated with site directed mutagenesis. Grh^Scr^ was made by randomizing the amino acids between 2-602 in Grh using the Peptide Nexus server (*77*). Codon composition was retained and the C-terminal amino acids 603-1032 were not modified. The structured minimal activation domain (Grh^173-228^) (*47*) was added back to Grh^Scr^, replacing amino acids 173-228 with the wild-type sequence to generate Grh^Scr+SD^. An N-terminal HA-tag was added to the cDNA encoding human GRHL2 (isoform 1). As a control, an N-terminal HA-tag was added to *Drosophila* Grh. Chimeric constructs were designed based on conserved regions defined by Traylor-Knowles et al. 2010 (*33*): HA-GRHL2.Grh (GRHL2^1-246^ fused to Grh^633-1032^) and HA-Grh.GRHL2 (Grh^1-632^ fused to GRHL2^247-624^). Stable cell lines were generated as described previously (*12*). S2 cells were cultured at 26°C in Schneider’s medium (Thermo Fisher Scientific) supplemented with 10% FBS (Omega Scientific) and antibiotic-antimycotic (Thermo Fisher Scientific). Cells were plated at 0.5 × 10^6^ cells per ml. After 24hrs, cells were transfected with 10 μg of plasmid DNA using Effectene transfection reagent (Qiagen). After an additional 24hrs, puromycin (Fisher Scientific) was at a final concentration of 2 μg/ml. Cells were recovered after 2-3 weeks of selection. Following recovery of stable cell lines, cells were cultured with 1μg/ml of puromycin.

#### Induction of TF expression

Cells were plated at 1 × 10^6^ cells per ml, and protein expression was induced by adding copper sulfate at the following concentrations: 100 µM Grh, 100 µM Grh^C800A,K807A^, 100 µM Grh^R806A^, 400 µM Grh^Scr^, 100 µM Grh^Scr+SD^, 200 µM HA-Grh, 450 µM HA-GRHL2, 450 µM HA-GRHL2.Grh, 450 µM HAGrh.GRHL2. Concentration of copper sulfate used for induction was determined based on approximation to physiological levels of expression as determined empirically by immunoblots (*12*). Cells were collected 48 hrs following induction for immunoblotting, ChIP-seq, or ATAC-seq.

#### Luciferase assays

The promoter of *hgo,* including the upstream Grh-binding site, was cloned into pGL3-Basic to drive Firefly luciferase expression (*68*). cDNAs were cloned into pAc5.1 for Grh protein expression. As previously published, transient transfections were performed in triplicate with 90 ng wild-type or mutant *hgo* reporter plasmid, 100 ng Grh-expression plasmid, 10 ng of actin-renilla plasmid and 100 ng empty expression plasmid using the Effectene Transfection Reagent (Qiagen) (*68*). Luciferase assays were performed on cell lysate using the Dual-luciferase assay kit (Promega). Fold activation was calculated relative to luciferase reads from controls transfected with 200 ng empty expression plasmid in place of the Grh-expression plasmid.

#### Immunoblotting

Proteins were separated by SDS-PAGE and transferred to a 0.45μm PVDF membrane at 4°C in transfer buffer (20% methanol, 25 mM Tris, 200 mM glycine) for 60min at 500mA. Membranes were blocked at room temperature in BLOTTO (2.5% nonfat dry milk, 0.5% BSA and 0.5% NP-40 in TBST) for 30 minutes at room temperature. Primary antibody incubations were performed overnight at 4°C at the following concentrations: anti- Grh (1:1,000) (*75*), anti-HA-peroxidase (1:500) (clone 3F10, Roche), anti-tubulin (1:10,000) (DM1A, Sigma). Secondary antibody incubation with goat anti-rabbit IgG-HRP (1:5,000) (BioRad) or goat anti-mouse IgG-HRP (1:6,000) (BioRad) was for 1 hr at room temperature. HRP activity was detected using SuperSignal West Pico PLUS chemiluminescent substrate (Thermo Fisher Scientific). Imaging was performed using film or an Azure Biosystems c600 imaging system.

#### ChIP sequencing

ChIP-seq was performed as described previously (*12*). Briefly, two replicates were collected with 25 × 10^6^ cells per replicate. Samples were fixed in 0.8% formaldehyde for 7 min and quenched with 125 mM glycine. Fixed chromatin was sonicated on a Covaris S220. IPs were incubated with 8uL anti-Grh (*75*) or 7.5 uL anti-HA (12CA5; Sigma-Aldrich) at 4°C overnight and purified using Dynabeads Protein A for Grh (20 uL; Thermo Fisher Scientific) or M-280 Sheep Anti-Mouse IgG for HA (60 uL; Thermo Fisher Scientific). The beads were washed, and chromatin was eluted. IPs and inputs were treated with RNaseA, and crosslinks were reversed. DNA was isolated by phenol:chloroform extraction and concentrated overnight by ethanol precipitation. Libraries were prepared using the NEB Next Ultra II kit. Grh-mutant libraries (Fig. 2) were sent to the Northwestern Sequencing Core (NUSeq) for sequence on the Illumina NextSeq 500 using 75bp single-end reads, additional reads were sequenced on the Illumina HiSeq 4000 using 50bp single-end reads. Grh^Scr^ libraries (Fig. 4) were sent to NUSeq for sequencing on the Illumina NovaSeq X Plus using 50bp paired-end reads. GRHL2 libraries (Fig. 5) were sent for sequencing at the UW Madison Biotechnology Center on the Illumina NovaSeq X Plus using 150bp paired-end reads.

#### ATAC sequencing

ATAC-seq was performed as described previously (*12*). For each experiment, two replicates were collected with 2 × 10^5^ cells per replicate. Cells were lysed in ATAC lysis buffer (10 mM Tris, 10 mM NaCl, 3 mM MgCl_2_, 0.1% NP-40), resuspended in buffer TD (Illumina), and tagmented with Tn5 enzyme (Illumina). DNA was purified with the MinElute Reaction Cleanup kit (Qiagen), and PCR amplified with IDT for Illumina Nextera DNA Unique Dual Indices in NEBNext Hi-Fi 2x PCR Master Mix. Libraries were purified with Axygen paramagentic beads (Thermo Fisher Scientific) at a ratio of 1.2x and eluted in 10 mM Tris. Samples were sequenced at the UW Madison Biotechnology Center on the Illumina NovaSeq X Plus using 150 bp paired-end reads.

#### Imaging mitotic retention

cDNAs encoding Grh, Grh^ΔN^, Grh^R806A^, or GRHL2 were cloned into pAc5.1 with an N-terminal GFP tag. Cells were plated in 35mm glass bottom dishes at 1.0 × 10^6^ cells/ml and incubated at 25°C for 30 minutes to allow for adherence. Cells were then transfected with 250 ng of plasmid encoding GFP-Grh and 250 ng of pAc-mCh-tubulin (DGRC stock# 1462) (*78*) using the Effectene transfection reagent (Qiagen). Cells were imaged 48 hrs after transfection on a Nikon Eclipse Ti2-E inverted fluorescent microscope. Mitotic cells were visually identified using mCh-tubulin to identify the spindle and then imaged through mitosis with a 60x objective every 10 seconds.

#### Drosophila culture and transgene expression

All flies were grown on molasses at 25°C unless otherwise noted. Transgenic flies were made by cloning cDNAs encoding Grh, Grh^ΔN^, Grh^R806A^, or GRHL2 into pUASt-attB with an N-terminal Halo tag and integrated at docking site VK14 on chromosome II using ФC31-mediated integration (BestGene, Chino Hills, CA). Transgenic expression was driven in a temperature-dependent manner. Fly lines carrying transgenes for Grh expression were crossed to *MS1096-GAL4,tubulin-GAL80^ts^*(*79*) and raised at 18°C. Larva were moved to 30°C for 24 hrs prior to dissection.

#### Salivary gland dissections and Halo ligand treatment

Salivary glands were dissected from third-instar larvae of the selected genotype in Schneider media supplemented with 5% fetal bovine serum and 1x Antibiotic-antimycotic. Glands were incubated for 15min with Halo ligands: 200 nM JF646 (Promega) for FRAP and imaging or 0.5 nM TMR (Promega) for single-molecule tracking. Following incubation, glands were washed three times for 10min each in media followed by a PBS rinse. Glands were then mounted onto polylysine-coated coverslips in Schneider media with 2.5% wt/vol methyl-cellulose as described previously (*80*).

#### Fluorescent recovery after photobleaching

Imaging of the salivary glands was performed on JF646 labelled salivary glands as described in DeHaro-Arbona et al. 2024. Individual nuclei were imaged with a 4.5× zoom, 512 × 512 pixel resolution, and settings were optimized for bleaching and scanning speed: pinhole was opened to 3.5-Airy, speed was 700 Hz. Effective bleaching was achieved by point bleaching. Images before and after bleaching were acquired every 0.4 s. After 50 images post bleaching, frame gap was increased to 1 s to minimize unintentional bleaching. FRAP curves were normalized as described in DeHaro-Arbona et al. His2Av-iRFP was imaged as a control.

#### Single molecule tracking

Single-molecule imaging was performed using a custom-built microscope as described previously (*80*, *81*). The TMR Halo ligand was excited at 561 nm under continuous illumination and emission was collected at 585 nm. Between 5 - 8 nuclei (Grh, 8; Grh^ΔN^, 6; Grh^R806A^, 5) were imaged with a 50ms exposure time for 5 minutes.

### Statistical analysis

#### ChIP sequencing analysis

Data was analyzed as previously described (*12*). Briefly, Reads were aligned to the *Drosophila melanogaster* genome (dm6) using Bowtie2 (v2.4.4). Unmapped reads, reads aligning to the mitochondrial genome or unplaced scaffolds, reads aligning more than once or low-quality reads (MAPQ<30) were removed. Peaks were called with MACS2 (v2.2) with the corresponding input as control, only peaks detected in both replicates were retained for downstream analyses. To generate bigWigs we used deeptools bamCoverage (v3.5.1) and z-score normalized across the genome.

#### ATAC sequencing analysis

Reads were processed as previously described (*12*). In brief, reads were trimmed for adapter sequences using NGmerge and aligned to the *Drosophila melanogaster* genome (dm6) using Bowtie2 (v2.4.4). Unmapped reads, reads aligning to the mitochondrial genome or unplaced scaffolds, reads aligning more than once or low-quality reads (MAPQ<30) were removed. Only fragments shorter than 100 bp were used for downstream analysis. Peaks were called with MACS2 (v2.2) and only peaks detected in both replicates were retained for downstream analysis. bigWig files were generated with deepTools bamCoverage (v3.5.1) and z-score normalized across the genome. Differential accessibility was determined using DESeq2 and compared induced samples to uninduced controls.

#### Single molecule tracking analysis

Analysis of single molecule trajectories was performed using a previously described pipeline (*81*). Briefly, single molecules were localized using a Gaussian fitting-based approach (*82*) and tracked using a multiple hypothesis tracking algorithm (*83*). Trajectories with a minimum of 4 time points were analyzed using a variational Bayesian treatment of Hidden Markov models (vbSPT) (*84*) and assigned to one of 3 states (D1, D2, or D3) defined by a unique Brownian motion diffusion coefficient.

#### Angle and anisotropy analyses

Anisotropy analyses were done as previously described except that only D3 (as identified by vbSPT) was used for analysis (*81*, *85*). The fold anisotropy metric f(180/0) was calculated as follows:

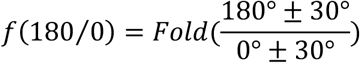

## Acknowledgments

We would like to thank the University of Wisconsin-Madison Biotechnology Center and the NUSeq Core Facility for sequencing. We acknowledge the Bloomington Drosophila Stock Center and the Drosophila Genome Resource Center for providing reagents and fly lines used in this study. MMF was supported by a T32GM007133. Experiments were supported by a NIH R35 GM136298 (MMH), a Wellcome Trust Investigator Award (212207/A/18) to SJB and a Royal Society International Exchange Award to SJB and MMH. MMH was also supported by a Vallee Scholar Award.

## Author contributions

MMF, FJA, SB, SJB and MMH designed the experiments. MMF, FJA, SB, AT, CR, AJR, AQR, RY and MMH performed the experiments and data analysis. MMF and MMH wrote the original draft. MMF, FJA, SB, PWL, SJB and MMH revised and edited the manuscript. MMF, MMH, and SJB acquired funding.

## Data and materials availability

ChIP-seq data is deposited to GEO accession number GSE315613. ATAC-seq data is deposited to GEO accession number GSE315642.

## REFERENCES

1. X.-Y. Li, S. Thomas, P. J. Sabo, M. B. Eisen, J. A. Stamatoyannopoulos, M. D. Biggin, The role of chromatin accessibility in directing the widespread, overlapping patterns of Drosophila transcription factor binding. Genome Biol 12, R34 (2011).

2. M. Iwafuchi-Doi, K. S. Zaret, Pioneer transcription factors in cell reprogramming. Genes Dev 28, 2679–2692 (2014).

3. A. Soufi, M. F. Garcia, A. Jaroszewicz, N. Osman, M. Pellegrini, K. S. Zaret, Pioneer transcription factors target partial DNA motifs on nucleosomes to initiate reprogramming. Cell 161, 555–568 (2015).

4. K. S. Zaret, Pioneer Transcription Factors Initiating Gene Network Changes. Annu. Rev. Genet. 54, 367–385 (2020).

5. E. D. Larson, A. J. Marsh, M. M. Harrison, Pioneering the developmental frontier. Molecular Cell 81, 1640–1650 (2021).

6. S. Stoeber, H. Godin, C. Xu, L. Bai, Pioneer factors: nature or nurture? Critical Reviews in Biochemistry and Molecular Biology, 1–15 (2024).

7. L. A. Cirillo, Binding of the winged-helix transcription factor HNF3 to a linker histone site on the nucleosome. The EMBO Journal 17, 244–254 (1998).

8. B. T. Donovan, H. Chen, P. Eek, Z. Meng, C. Jipa, S. Tan, L. Bai, M. G. Poirier, Basic helix-loop-helix pioneer factors interact with the histone octamer to invade nucleosomes and generate nucleosome-depleted regions. Molecular Cell 83, 1251–1263.e6 (2023).

9. M. Fernandez Garcia, C. D. Moore, K. N. Schulz, O. Alberto, G. Donague, M. M. Harrison, H. Zhu, K. S. Zaret, Structural Features of Transcription Factors Associating with Nucleosome Binding. Molecular Cell 75, 921–932.e6 (2019).

10. J. Gassler, W. Kobayashi, I. Gáspár, S. Ruangroengkulrith, A. Mohanan, L. Gómez Hernández, P. Kravchenko, M. Kümmecke, A. Lalic, N. Rifel, R. J. Ashburn, M. Zaczek, A. Vallot, L. Cuenca Rico, S. Ladstätter, K. Tachibana, Zygotic genome activation by the totipotency pioneer factor Nr5a2. Science 378, 1305–1315 (2022).

11. S. L. McDaniel, T. J. Gibson, K. N. Schulz, M. Fernandez Garcia, M. Nevil, S. U. Jain, P. W. Lewis, K. S. Zaret, M. M. Harrison, Continued Activity of the Pioneer Factor Zelda Is Required to Drive Zygotic Genome Activation. Molecular Cell 74, 185–195.e4 (2019).

12. T. J. Gibson, E. D. Larson, M. M. Harrison, Protein-intrinsic properties and context-dependent effects regulate pioneer factor binding and function. Nat Struct Mol Biol 31, 548–558 (2024).

13. J. Lerner, A. Katznelson, J. Zhang, K. S. Zaret, Different chromatin-scanning modes lead to targeting of compacted chromatin by pioneer factors FOXA1 and SOX2. Cell Reports 42, 112748 (2023).

14. S. Sakong, B. Fierz, D. Suter, Electrostatic properties of disordered regions control transcription factor search and pioneer activity. bioRxiv [Preprint] (2025). 10.1101/2025.03.07.641980.

15. S. Brodsky, T. Jana, K. Mittelman, M. Chapal, D. K. Kumar, M. Carmi, N. Barkai, Intrinsically Disordered Regions Direct Transcription Factor *In Vivo* Binding Specificity. Molecular Cell 79, 459–471.e4 (2020).

16. D. A. Garcia, T. A. Johnson, D. M. Presman, G. Fettweis, K. Wagh, L. Rinaldi, D. A. Stavreva, V. Paakinaho, R. A. M. Jensen, S. Mandrup, A. Upadhyaya, G. L. Hager, An intrinsically disordered region-mediated confinement state contributes to the dynamics and function of transcription factors. Mol Cell 81, 1484–1498.e6 (2021).

17. F. Jonas, Y. Navon, N. Barkai, Intrinsically disordered regions as facilitators of the transcription factor target search. Nat Rev Genet 26, 424–435 (2025).

18. X. A. Feng, M. Yamadi, Y. Fu, K. M. Ness, C. Liu, I. Ahmed, G. D. Bowman, M. E. Johnson, T. Ha, C. Wu, GAGA zinc finger transcription factor searches chromatin by 1D–3D facilitated diffusion. Nat Struct Mol Biol 32, 2359–2370 (2025).

19. X. Tang, T. Li, S. Liu, J. Wisniewski, Q. Zheng, Y. Rong, L. D. Lavis, C. Wu, Kinetic principles underlying pioneer function of GAGA transcription factor in live cells. Nat Struct Mol Biol 29, 665–676 (2022).

20. M. Már, K. Nitsenko, P. O. Heidarsson, Multifunctional Intrinsically Disordered Regions in Transcription Factors. Chemistry – A European Journal 29, e202203369 (2023).

21. Y. Chen, C. Cattoglio, G. M. Dailey, Q. Zhu, R. Tjian, X. Darzacq, Mechanisms governing target search and binding dynamics of hypoxia-inducible factors. Elife 11, e75064 (2022).

22. M. Iwafuchi, I. Cuesta, G. Donahue, N. Takenaka, A. B. Osipovich, M. A. Magnuson, H. Roder, S. H. Seeholzer, P. Santisteban, K. S. Zaret, Gene network transitions in embryos depend upon interactions between a pioneer transcription factor and core histones. Nat Genet 52, 418–427 (2020).

23. Z. Wang, B. Wang, D. Niu, C. Yin, Y. Bi, C. Cattoglio, K. M. Loh, L. D. Lavis, H. Ge, W. Deng, Mesoscale chromatin confinement facilitates target search of pioneer transcription factors in live cells. Nat Struct Mol Biol 32, 125–136 (2025).

24. M. V. Staller, Transcription factors perform a 2-step search of the nucleus. Genetics 222, iyac111 (2022).

25. B. R. Sabari, A. Dall’Agnese, A. Boija, I. A. Klein, E. L. Coffey, K. Shrinivas, B. J. Abraham, N. M. Hannett, A. V. Zamudio, J. C. Manteiga, C. H. Li, Y. E. Guo, D. S. Day, J. Schuijers, E. Vasile, S. Malik, D. Hnisz, T. I. Lee, I. I. Cisse, R. G. Roeder, P. A. Sharp, A. K. Chakraborty, R. A. Young, Coactivator condensation at super-enhancers links phase separation and gene control. Science 361, eaar3958 (2018).

26. J. J. Ferrie, J. P. Karr, R. Tjian, X. Darzacq, “Structure”-function relationships in eukaryotic transcription factors: The role of intrinsically disordered regions in gene regulation. Molecular Cell 82, 3970–3984 (2022).

27. S. Chong, C. Dugast-Darzacq, Z. Liu, P. Dong, G. M. Dailey, C. Cattoglio, A. Heckert, S. Banala, L. Lavis, X. Darzacq, R. Tjian, Imaging dynamic and selective low-complexity domain interactions that control gene transcription. Science 361, eaar2555 (2018).

28. S. J. Bray, F. C. Kafatos, Developmental function of Elf-1: an essential transcription factor during embryogenesis in Drosophila. Genes Dev. 5, 1672–1683 (1991).

29. A. E. Uv, E. J. Harrison, S. J. Bray, Tissue-specific splicing and functions of the Drosophila transcription factor Grainyhead. Mol Cell Biol 17, 6727–6735 (1997).

30. A. E. Uv, C. R. Thompson, S. J. Bray, The Drosophila tissue-specific factor Grainyhead contains novel DNA-binding and dimerization domains which are conserved in the human protein CP2. 12.

31. K. Venkatesan, H. R. McManus, C. C. Mello, T. F. Smith, U. Hansen, Functional conservation between members of an ancient duplicated transcription factor family, LSF/Grainyhead. Nucleic Acids Res 31, 4304–4316 (2003).

32. T. Wilanowski, A. Tuckfield, L. Cerruti, S. O’Connell, R. Saint, V. Parekh, J. Tao, J. M. Cunningham, S. M. Jane, A highly conserved novel family of mammalian developmental transcription factors related to Drosophila grainyhead. Mechanisms of Development 114, 37–50 (2002).

33. N. Traylor-Knowles, U. Hansen, T. Q. Dubuc, M. Q. Martindale, L. Kaufman, J. R. Finnerty, The evolutionary diversification of LSF and Grainyhead transcription factors preceded the radiation of basal animal lineages. BMC Evolutionary Biology 10, 101 (2010).

34. R. M. Reese, M. M. Harrison, E. T. Alarid, Grainyhead-like Protein 2: The Emerging Role in Hormone-Dependent Cancers and Epigenetics. Endocrinology 160, 1275–1288 (2019).

35. A. Paré, M. Kim, M. T. Juarez, S. Brody, W. McGinnis, The functions of grainy head-like proteins in animals and fungi and the evolution of apical extracellular barriers. PLoS One 7, e36254 (2012).

36. K. A. Mace, J. C. Pearson, W. McGinnis, An Epidermal Barrier Wound Repair Pathway in Drosophila Is Mediated by grainy head. Science 308, 381–385 (2005).

37. J. Hemphälä, A. Uv, R. Cantera, S. Bray, C. Samakovlis, Grainy head controls apical membrane growth and tube elongation in response to Branchless/FGF signalling. Development 130, 249–258 (2003).

38. C. Nüsslein-Volhard, E. Wieschaus, H. Kluding, Mutations affecting the pattern of the larval cuticle inDrosophila melanogaster. Wilhelm Roux’ Archiv 193, 267–282 (1984).

39. S. J. Bray, W. A. Johnson, J. Hirsh, U. Heberlein, R. Tjian, A cis-acting element and associated binding factor required for CNS expression of the Drosophila melanogaster dopa decarboxylase gene. The EMBO Journal 7, 177–188 (1988).

40. B. D. Dynlacht, L. D. Attardi, A. Admon, M. Freeman, R. Tjian, Functional analysis of NTF-1, a developmentally regulated Drosophila transcription factor that binds neuronal cis elements. Genes Dev. 3, 1677–1688 (1989).

41. S. B. Ting, T. Wilanowski, L. Cerruti, L.-L. Zhao, J. M. Cunningham, S. M. Jane, The identification and characterization of human Sister-of-Mammalian Grainyhead (SOM) expands the grainyhead-like family of developmental transcription factors. Biochem. J. 370, 953–962 (2003).

42. A. Auden, J. Caddy, T. Wilanowski, S. B. Ting, J. M. Cunningham, S. M. Jane, Spatial and temporal expression of the *Grainyhead*-like transcription factor family during murine development. Gene Expression Patterns 6, 964–970 (2006).

43. S. Riethdorf, S. Frey, S. Santjer, M. Stoupiec, B. Otto, L. Riethdorf, C. Koop, W. Wilczak, R. Simon, G. Sauter, K. Pantel, V. Assmann, Diverse expression patterns of the EMT suppressor grainyhead-like 2 (GRHL2) in normal and tumour tissues. International Journal of Cancer 138, 949–963 (2016).

44. J. Jacobs, M. Atkins, K. Davie, H. Imrichova, L. Romanelli, V. Christiaens, G. Hulselmans, D. Potier, J. Wouters, I. I. Taskiran, G. Paciello, C. B. González-Blas, D. Koldere, S. Aibar, G. Halder, S. Aerts, The transcription factor Grainy head primes epithelial enhancers for spatiotemporal activation by displacing nucleosomes. Nat Genet 50, 1011–1020 (2018).

45. M. Nevil, T. J. Gibson, C. Bartolutti, A. Iyengar, M. M. Harrison, Establishment of chromatin accessibility by the conserved transcription factor Grainy head is developmentally regulated. Development, dev.185009 (2020).

46. A. F. Chen, A. J. Liu, R. Krishnakumar, J. W. Freimer, B. DeVeale, R. Blelloch, GRHL2-Dependent Enhancer Switching Maintains a Pluripotent Stem Cell Transcriptional Subnetwork after Exit from Naive Pluripotency. Cell Stem Cell 23, 226–238.e4 (2018).

47. L. D. Attardi, R. Tjian, Drosophila tissue-specific transcription factor NTF-1 contains a novel isoleucine-rich activation motif. Genes & Development 7, 1341–1353 (1993).

48. H. A. Ertl, E. X. Bayala, M. A. Siddiq, P. J. Wittkopp, Divergence of Grainy head affects chromatin accessibility, gene expression, and embryonic viability in Drosophila melanogaster. bioRxiv, 2024.04.07.588430 (2024).

49. F. Zhu, L. Farnung, E. Kaasinen, B. Sahu, Y. Yin, B. Wei, S. O. Dodonova, K. R. Nitta, E. Morgunova, M. Taipale, P. Cramer, J. Taipale, The interaction landscape between transcription factors and the nucleosome. Nature 562, 76–81 (2018).

50. V. Ramalingam, X. Yu, B. D. Slaughter, J. R. Unruh, K. J. Brennan, A. Onyshchenko, J. J. Lange, M. Natarajan, M. Buck, J. Zeitlinger, Lola-I is a promoter pioneer factor that establishes de novo Pol II pausing during development. Nat Commun 14, 5862 (2023).

51. P. T. Lowary, J. Widom, New DNA sequence rules for high affinity binding to histone octamer and sequence-directed nucleosome positioning1. Journal of Molecular Biology 276, 19–42 (1998).

52. R. S. Isaac, F. Jiang, J. A. Doudna, W. A. Lim, G. J. Narlikar, R. Almeida, Nucleosome breathing and remodeling constrain CRISPR-Cas9 function. eLife 5, e13450 (2016).

53. M. G. Poirier, M. Bussiek, J. Langowski, J. Widom, Spontaneous Access to DNA Target Sites in Folded Chromatin Fibers. Journal of Molecular Biology 379, 772–786 (2008).

54. W. Antonin, H. Neumann, Chromosome condensation and decondensation during mitosis. Current Opinion in Cell Biology 40, 15–22 (2016).

55. J. Gottesfeld, Mitotic repression of the transcriptional machinery. Trends in Biochemical Sciences 22, 197–202 (1997).

56. N. Naumova, M. Imakaev, G. Fudenberg, Y. Zhan, B. R. Lajoie, L. A. Mirny, J. Dekker, Organization of the Mitotic Chromosome. Science 342, 948–953 (2013).

57. M. Bellec, J. Dufourt, G. Hunt, H. Lenden-Hasse, A. Trullo, A. Zine El Aabidine, M. Lamarque, M. M. Gaskill, H. Faure-Gautron, M. Mannervik, M. M. Harrison, J.-C. Andrau, C. Favard, O. Radulescu, M. Lagha, The control of transcriptional memory by stable mitotic bookmarking. Nat Commun 13, 1176 (2022).

58. A. Chervova, A. Molliex, H. I. Baymaz, T. Papadopoulou, F. Mueller, E. Hercul, D. Fournier, A. Dubois, N. Gaiani, P. Beli, N. Festuccia, P. Navarro, Mitotic bookmarking redundancy by nuclear receptors mediates robust post-mitotic reactivation of the pluripotency network. [Preprint] (2022). 10.1101/2022.11.28.518105.

59. C. Deluz, E. T. Friman, D. Strebinger, A. Benke, M. Raccaud, A. Callegari, M. Leleu, S. Manley, D. M. Suter, A role for mitotic bookmarking of SOX2 in pluripotency and differentiation. Genes Dev. 30, 2538–2550 (2016).

60. X. Liu, J. Shen, L. Xie, Z. Wei, C. Wong, Y. Li, X. Zheng, P. Li, Y. Song, Mitotic Implantation of the Transcription Factor Prospero via Phase Separation Drives Terminal Neuronal Differentiation. Developmental Cell 52, 277–293.e8 (2020).

61. R. Silvério-Alves, I. Kurochkin, A. Rydström, C. Vazquez Echegaray, J. Haider, M. Nicholls, C. Rode, L. Thelaus, A. Y. Lindgren, A. G. Ferreira, R. Brandão, J. Larsson, M. F. T. R. De Bruijn, J. Martin-Gonzalez, C.-F. Pereira, GATA2 mitotic bookmarking is required for definitive haematopoiesis. Nat Commun 14, 4645 (2023).

62. N. Festuccia, A. Dubois, S. Vandormael-Pournin, E. Gallego Tejeda, A. Mouren, S. Bessonnard, F. Mueller, C. Proux, M. Cohen-Tannoudji, P. Navarro, Mitotic binding of Esrrb marks key regulatory regions of the pluripotency network. Nat Cell Biol 18, 1139–1148 (2016).

63. J. M. Caravaca, G. Donahue, J. S. Becker, X. He, C. Vinson, K. S. Zaret, Bookmarking by specific and nonspecific binding of FoxA1 pioneer factor to mitotic chromosomes. Genes & Development 27, 251–260 (2013).

64. Y. Liu, B. Pelham-Webb, D. C. Di Giammartino, J. Li, D. Kim, K. Kita, N. Saiz, V. Garg, A. Doane, P. Giannakakou, A.-K. Hadjantonakis, O. Elemento, E. Apostolou, Widespread Mitotic Bookmarking by Histone Marks and Transcription Factors in Pluripotent Stem Cells. Cell Rep 19, 1283–1293 (2017).

65. R. M. Price, M. A. Budzyński, J. Shen, J. E. Mitchell, J. Z. J. Kwan, S. S. Teves, Heat shock transcription factors demonstrate a distinct mode of interaction with mitotic chromosomes. Nucleic Acids Research 51, 5040–5055 (2023).

66. S. S. Teves, L. An, A. S. Hansen, L. Xie, X. Darzacq, R. Tjian, A dynamic mode of mitotic bookmarking by transcription factors. eLife 5, e22280 (2016).

67. Q. Ming, Y. Roske, A. Schuetz, K. Walentin, I. Ibraimi, K. M. Schmidt-Ott, U. Heinemann, Structural basis of gene regulation by the Grainyhead/CP2 transcription factor family. Nucleic Acids Research 46, 2082–2095 (2018).

68. M. Nevil, E. R. Bondra, K. N. Schulz, T. Kaplan, M. M. Harrison, Stable Binding of the Conserved Transcription Factor Grainy Head to its Target Genes Throughout *Drosophila melanogaster* Development. Genetics 205, 605–620 (2017).

69. J. T. Lis, Imaging Drosophila gene activation and polymerase pausing in vivo. Nature 450, 198–202 (2007).

70. J. Liu, N. B. Perumal, C. J. Oldfield, E. W. Su, V. N. Uversky, A. K. Dunker, Intrinsic disorder in transcription factors. Biochemistry 45, 6873–6888 (2006).

71. J. Dufourt, A. Trullo, J. Hunter, C. Fernandez, J. Lazaro, M. Dejean, L. Morales, S. Nait-Amer, K. N. Schulz, M. M. Harrison, C. Favard, O. Radulescu, M. Lagha, Temporal control of gene expression by the pioneer factor Zelda through transient interactions in hubs. Nat Commun 9, 5194 (2018).

72. M. A. F. Soares, D. S. Soares, V. Teixeira, A. Heskol, R. B. Bressan, S. M. Pollard, R. A. Oliveira, D. S. Castro, Hierarchical reactivation of transcription during mitosis-to-G1 transition by Brn2 and Ascl1 in neural stem cells. Genes Dev. 35, 1020–1034 (2021).

73. A. L. Sanborn, B. T. Yeh, J. T. Feigerle, C. V. Hao, R. J. Townshend, E. Lieberman Aiden, R. O. Dror, R. D. Kornberg, Simple biochemical features underlie transcriptional activation domain diversity and dynamic, fuzzy binding to Mediator. eLife 10, e68068 (2021).

74. A. J. Ruthenburg, H. Li, T. A. Milne, S. Dewell, R. K. McGinty, M. Yuen, B. Ueberheide, Y. Dou, T. W. Muir, D. J. Patel, C. D. Allis, Recognition of a Mononucleosomal Histone Modification Pattern by BPTF via Multivalent Interactions. Cell 145, 692–706 (2011).

75. M. M. Harrison, M. R. Botchan, T. W. Cline, Grainyhead and Zelda compete for binding to the promoters of the earliest-expressed Drosophila genes. Developmental Biology 345, 248–255 (2010).

76. D. C. Hamm, E. R. Bondra, M. M. Harrison, Transcriptional Activation Is a Conserved Feature of the Early Embryonic Factor Zelda That Requires a Cluster of Four Zinc Fingers for DNA Binding and a Low-complexity Activation Domain*. Journal of Biological Chemistry 290, 3508–3518 (2015).

77. Scramble Peptide or Protein Sequence. https://peptidenexus.com/article/sequence-scrambler.

78. R. D. Vale, J. A. Spudich, E. R. Griffis, Dynamics of myosin, microtubules, and Kinesin-6 at the cortex during cytokinesis in Drosophila S2 cells. J Cell Biol 186, 727–738 (2009).

79. F. J. DeHaro-Arbona, C. Roussos, S. Baloul, J. Townson, M. J. Gómez Lamarca, S. Bray, Dynamic modes of Notch transcription hubs conferring memory and stochastic activation revealed by live imaging the co-activator Mastermind. Elife 12, RP92083 (2024).

80. M. J. Gomez-Lamarca, J. Falo-Sanjuan, R. Stojnic, S. Abdul Rehman, L. Muresan, M. L. Jones, Z. Pillidge, G. Cerda-Moya, Z. Yuan, S. Baloul, P. Valenti, K. Bystricky, F. Payre, K. O’Holleran, R. Kovall, S. J. Bray, Activation of the Notch Signaling Pathway In Vivo Elicits Changes in CSL Nuclear Dynamics. Dev Cell 44, 611–623.e7 (2018).

81. S. Baloul, C. Roussos, M. Gomez-Lamarca, L. Muresan, S. Bray, Changes in searching behaviour of CSL transcription complexes in Notch active conditions. Life Sci Alliance 7, e202302336 (2023).

82. M. Ovesný, P. Křížek, J. Borkovec, Z. Švindrych, G. M. Hagen, ThunderSTORM: a comprehensive ImageJ plug-in for PALM and STORM data analysis and super-resolution imaging. Bioinformatics 30, 2389–2390 (2014).

83. N. Chenouard, I. Bloch, J.-C. Olivo-Marin, Multiple Hypothesis Tracking for Cluttered Biological Image Sequences. IEEE Transactions on Pattern Analysis and Machine Intelligence 35, 2736–3750 (2013).

84. F. Persson, M. Lindén, C. Unoson, J. Elf, Extracting intracellular diffusive states and transition rates from single-molecule tracking data. Nat Methods 10, 265–269 (2013).

85. A. S. Hansen, A. Amitai, C. Cattoglio, R. Tjian, X. Darzacq, Guided nuclear exploration increases CTCF target search efficiency. Nat Chem Biol 16, 257–266 (2020).

86. R. J. Emenecker, D. Griffith, A. S. Holehouse, Metapredict: a fast, accurate, and easy-to-use predictor of consensus disorder and structure. Biophysical Journal 120, 4312–4319 (2021).

87. A. S. Holehouse, R. K. Das, J. N. Ahad, M. O. G. Richardson, R. V. Pappu, CIDER: Resources to Analyze Sequence-Ensemble Relationships of Intrinsically Disordered Proteins. Biophys J 112, 16–21 (2017).

88. P. Machanick, T. L. Bailey, MEME-ChIP: motif analysis of large DNA datasets. Bioinformatics 27, 1696–1697 (2011).

