## Supplemental Figures for "Pioneer-factor activity requires stable chromatin occupancy mediated by both sequence-specific binding and disordered protein domains"

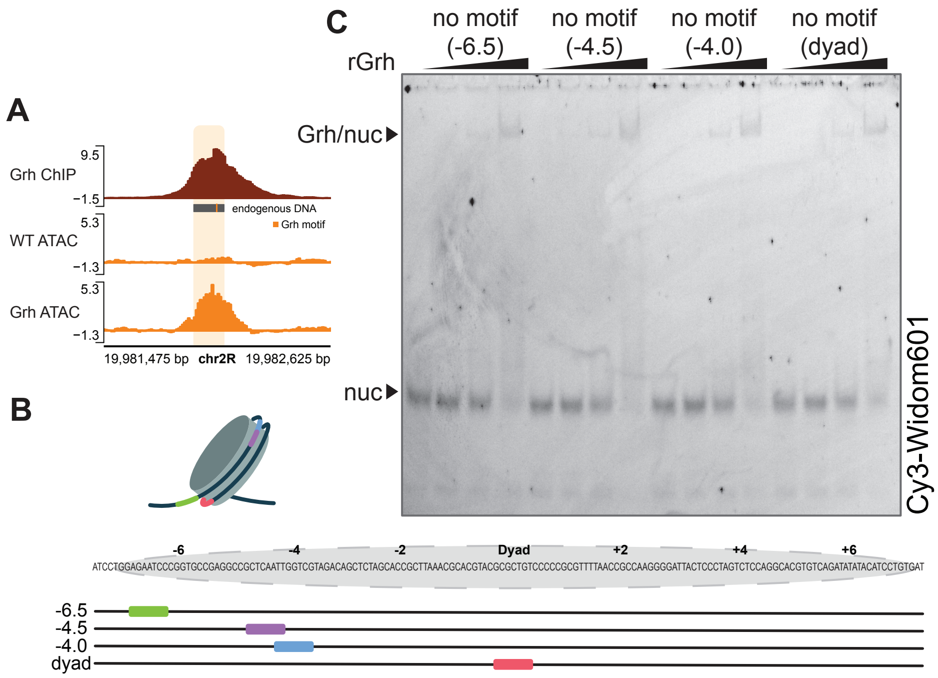


**Figure S1. Nucleosomal DNA sequences** (A) Genome browser tracks of the endogenous region used to make mononucleosomes. ATAC-seq data before induction (WT ATAC) and upon expression of Grh (Grh ATAC) demonstrate the increase in chromatin accessibility upon Grh expression. Grh ChIP-seq data upon expression of Grh (Grh ChIP). sequence used for the endogenous DNA is marked in grey with the Grh motif highlighted in orange. (B) Widom 601 sequence with Grh motifs marked at the noted positions. (C) EMSA with increasing concentrations of recombinant Grh (rGrh) (0 - 25nM) incubated with Cy3-labeled Widom 601 nucleosomes without the Grh motif. Same gel as Figure 1B but imaged for Cy3.

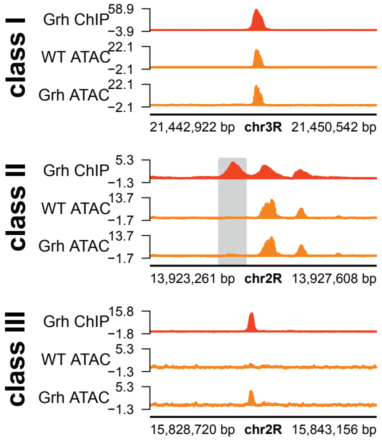


**Figure S2. Grh binds and opens chromatin in S2 cells.** Genome browser tracks showing ChIP-seq signal in cells expressing ectopic Grh and ATAC-seq signal before (WT ATAC) and after (Grh ATAC) induction of Grh. Example track shown of all three classes. Class I: accessible and bound by Grh. Class II: inaccessible and bound by Grh. Class III: inaccessible, bound and opened by Grh (as shown in Gibson et al. 2024 (*12*)).

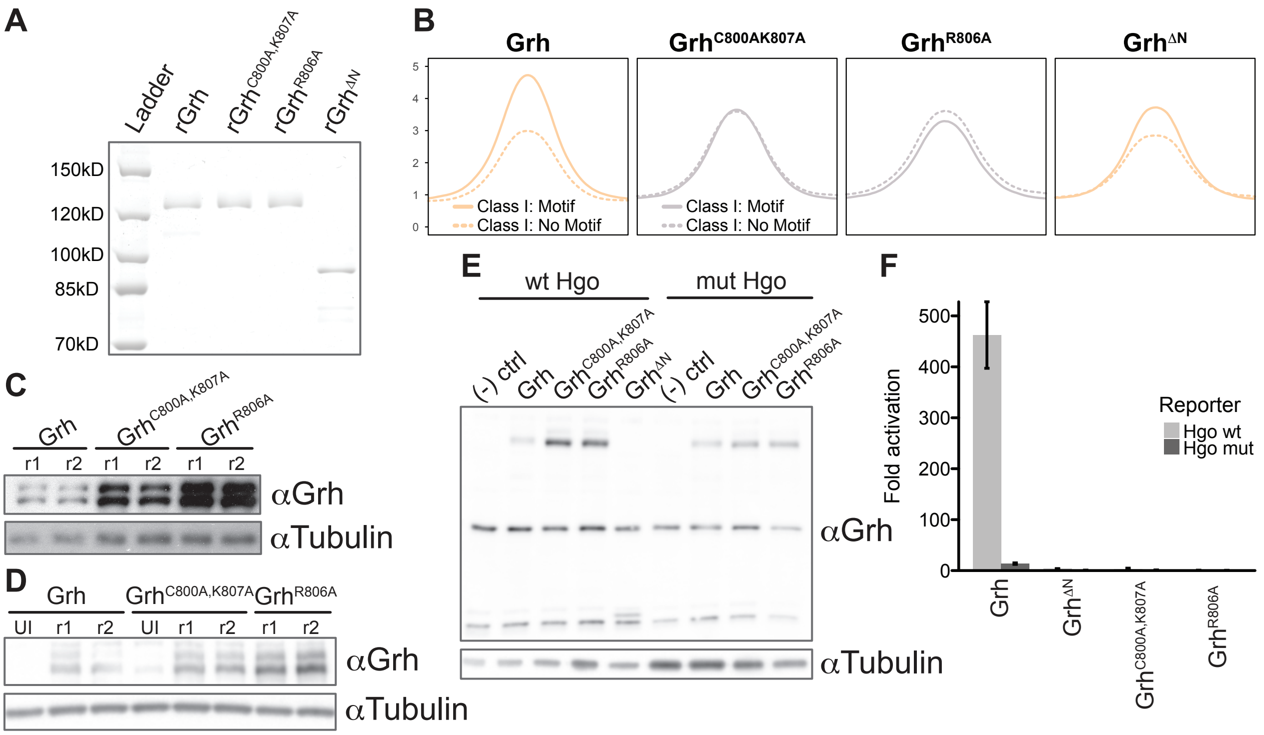


**Figure S3. Grh requires the DBD and regions outside for activating gene expression in a motif-specific way.** (A) Coomassie stain of an SDS-PAGE gel of recombinant Grh proteins: wild-type Grh, $Grh^{C800A,K807A}$, $Grh^{R806A}$ and $Grh^{\Delta N}$. (B) Metaplots showing Grh binding (ChIP-seq) at Class I sites containing the canonical Grh motif or without the motif (no motif). (C) Western blot showing expression of wild-type Grh, $Grh^{C800A,K807A}$ and $Grh^{R806A}$ in S2 cells used for ChIP-seq in Figure 2E. (D) Western blot showing expression of wild-type Grh, $Grh^{C800A,K807A}$ and $Grh^{R806A}$ in S2 cells used for ATAC-seq in Figure 2D. UI = uninduced, r1= replicate one, r2 = replicate two. (E) Western blot showing expression levels of wild-type Grh, $Grh^{C800A,K807A}$, $Grh^{R806A}$ and $Grh^{\Delta N}$ in S2 cells used for luciferase assay in F. (F) Fold activation of the wild-type or mutant *hgo* luciferase reporter by either wild-type Grh, $Grh^{C800A,K807A}$, $Grh^{R806A}$ or $Grh^{\Delta N}$. n = 3 technical replicates.

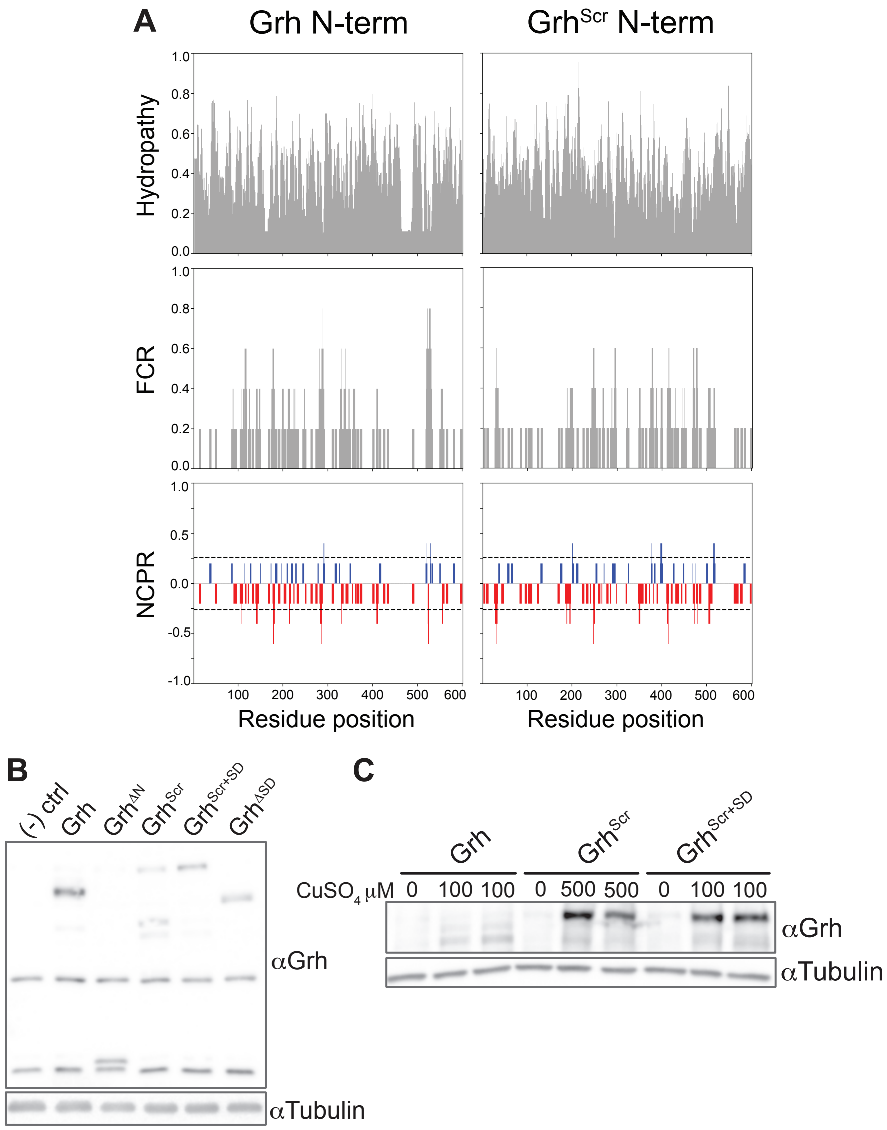


**Figure S4. The intrinsically disordered N-terminus can contribute to pioneering independed of amino acid sequence.** (A) Mean hydropathy, fraction of changed residues (FCR) and net charge per residue (NCPR) was calculated for groups of 5 residues (blob 5) with CIDER (87) (B) Western blot showing expression levels of wild-type Grh, $Grh^{\Delta N}$, $Grh^{Scr}$, $Grh^{Scr+SD}$ and $Grh^{\Delta SD}$ in S2 cells used for luciferase assay in Figure 4B. (B) Western blot showing expression of wild-type Grh, $Grh^{Scr}$ and $Grh^{Scr+SD}$ in S2 cells used for ChIP-seq and ATAC-seq in Figure 4C and 4D.

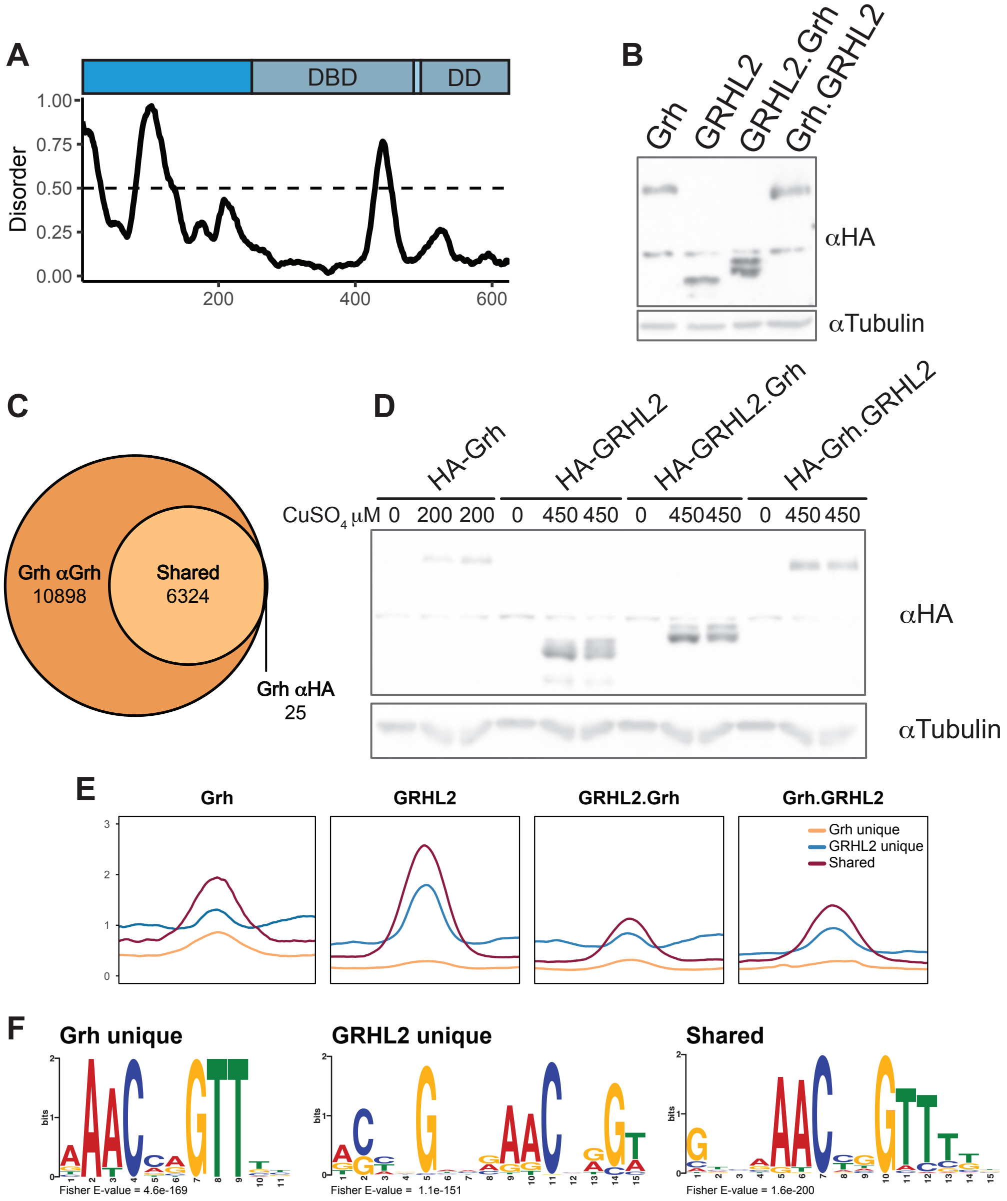


**Figure S5. Diverse N-terminal domains are sufficient for pioneering.** (A) Graph of predicted disorder score (from Metapredict (86)) for GRHL2. (B). Western blot showing expression levels of *Drosophila* Grh, human GRHL2, or chimeras (GRHL2.Grh, Grh.GRHL2) in S2 cells used for luciferase assay in Figure 5A. (C) Overlap between peaks called using the αGrh antibody or αHA antibody. (D) Western blot showing expression of *Drosophila* Grh, human GRHL2, or chimeras (GRHL2.Grh, Grh.GRHL2) in S2 cells used for ChIP-seq and ATAC-seq in Figure 5B and C. (E) Metaplots showing enrichment of binding at inaccessible sites unique to Grh or GRHL2 and shared sites. (F) Top motif enrichment for inaccessible sites uniquely bound by Grh or GRHL2 and shared sites identified by MEME-suite (88).

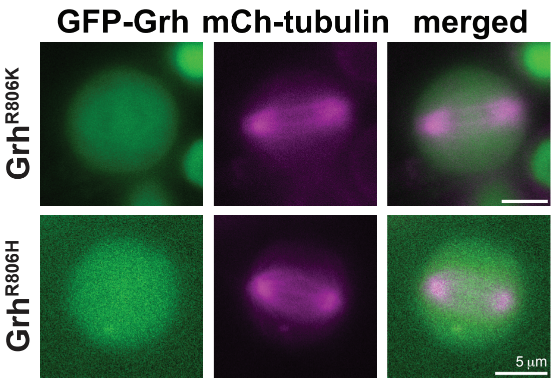


**Figure S6. Disruption of charge is not resposible for loss of Grh retention.** Images of GFP-tagged Grh proteins (green) and mCherry-tubulin (magenta) at anaphase in S2 cells.
